# Deciphering Cellular Ecosystems Driving Tumor Progression and Immune Escape from Spatial Transcriptomics and Single-Cell with COMPOTES

**DOI:** 10.1101/2025.10.27.684744

**Authors:** Loïc Herpin, Anaïs Chossegros, Roberta Codato, Josep Monserrat Sanchez, Jean El Khoury, Simon Grouard, Valérie Ducret, Alex Cornish, Baptiste Gross, the MOSAIC Consortium, Elodie Pronier, Caroline Hoffmann, Alberto Romagnoni, Eric Durand, Almudena Espin Perez, Quentin Bayard

## Abstract

Cell-cell communication is central to understanding the complex interactions within the tumor microenvironment. However, current methods fail to identify recurrent communication patterns across patient cohorts from spatial transcriptomics, as they are often limited to single samples or lack essential spatial context. Yet this is essential for understanding how local environments influence cell phenotype and states, and shape the entire cellular ecosystem.

We introduce a machine-learning approach that models local, spatially aware ligand-receptor interactions and uses matrix factorization to extract global multicellular programs from large cohorts representing the complex biology of cancer.

Applied to a multimodal muscle-invasive bladder cancer cohort of 146 patients, it uncovered 45 communication programs defined by distinct ligand-receptor pairs and cellular niches. In particular, we identified a conserved immune program linked to stalled anti-tumor immunity and a program linking *KMT2D* loss-of-function mutations with early-stage (T2) tumors, intense proliferation and a favorable response to neoadjuvant chemotherapy.

## Introduction

Cell-cell communication (CCC) is fundamental to cellular behavior within the Tumor Micro-Environment (TME), governing processes like cell survival, proliferation, and immune evasion. These interactions, mediated by ligands and receptors, are context-dependent, with crosstalk between cancer, immune, and stromal cells shaping tumor progression^1,2^. Consequently, disrupted signaling pathways can drive malignancy, making them prime therapeutic targets^3–5^.

Direct exploration of intercellular interactions is a difficult process. Over the last decade, the emergence of single-cell transcriptomics (scRNA-seq) and spatialized transcriptomics sequencing technologies has opened an alternative way to explore the multicellular environment by using the spatial context and transcriptomics data at the granular resolution of individual cells. These technologies have been widely used to infer cell-cell interactions^6–10^.

Many CCC inference methods follow a common framework, quantifying ligand-receptor interactions (LRIs) using expression data and curated databases^6,7,11–13^. These databases vary in complexity; for instance, CellPhoneDB^8^ accounts for heteromeric protein complexes, while CellChatDB^9^ incorporates the detailed structural composition of signaling molecules, including co-receptors and multimeric complexes.

To derive LRI scores, most methods apply standard statistical measures including mean, geometric mean, and product of ligand/receptor expression values, which can then be used to filter interactions based on specific thresholds^8,14,15,16^. Some use more complex scoring, such as differential expression analysis^7,8^ to retrieve ligand and receptor (LR) pairs with significant expression change. Others model the physical properties of molecules for a finer representation of the TME^14^. Finally, some methods determine significant interactions by using permutation tests or machine learning methods ^8,9,17,18^. A cell-cell interaction score can then be derived from all these possibly interacting pairs of cells. As opposed to typical agglomerative methods which will only consider pairs of ligands receptors as directly connected to clusters of cells, some methods try to provide deeper insight into interactions, heterogeneity and specific subpopulations of cells^19,20^,using graph representation ^20^or focusing on intercellular interactions. ^6,11,21,22^.

However, CCC inference from scRNA-seq has key limitations^23^: the lack of spatial information can generate false positives, while averaging signal at pre-defined cell-type levels often obscures communication events between distinct cellular subpopulations or cell states^6,12^. The absence of a ground truth for downstream signaling further complicates validation.

The advent of spatial transcriptomics has mitigated the spatial issue^11–13,15,24–31^, enabling the constraint of interactions by cellular proximity and sparking new modeling approaches that account for distance^22,24,27,30,32^ or competition between LR pairs^27^. Concurrently, other methods have improved CCC inference by incorporating intracellular signaling pathways^22,33–36^ or by scaling the analysis to multiple samples^29,37–39^. These multi-sample approaches, often employing matrix factorization^38–41^, are crucial for identifying recurrent communication programs associated with disease progression or drug resistance. Despite these parallel advances, to our knowledge, no work has yet integrated the local, detailed modeling of LRI from spatial transcriptomics with the global extraction of high-level communication patterns across multiple samples.

Here, we present COMPOTES (cell-cell COMmunication Programs Of the Tumor microEnvironment across Spatial transcriptomic samples), a method to extract multicellular communication programs from multiple spatial transcriptomics samples by modeling LRIs based on the product of expression of the gene coding for these proteins, modeling distance, protein complexes, and ligand competition. Applying our method to the Muscle-Invasive Bladder Cancer (MIBC) subset of the MOSAIC dataset^42^, we discovered 45 communication programs. They revealed granular communication processes at multiple scales, some associated with clinical variables such as response to treatment or stages and tumor subtypes, and many of which generalized across other cancer types.

Our framework validates known biological paradigms, such as identifying a conserved immune program that captures the known paradox of an inflamed yet immunosuppressed TME, characterized by a stalled anti-tumor response, T-cell exhaustion, and regulatory immune cell infiltration. It also uncovers a novel, MIBC-specific communication program that defines a paradoxical state of “high-proliferation, high-vulnerability.” This program, linked to inactivating KMT2D mutations, is most active in early-stage (T2) tumors, and its intense proliferation explains a favorable response to neoadjuvant chemotherapy. Deployed by distinct tumor subclones, it contains intrinsic molecular brakes that physically constrain its expansion to the tumor core. Thus, our approach reveals a previously uncharacterized, epigenetically-driven program whose inherent fragility represents a barrier to progression and a key therapeutic vulnerability, offering new insights into the early-stage MIBC dynamics.

## Results

### COMPOTES: an hybrid method for multiscale CCC analysis

We introduce COMPOTES, the first method for extracting latent multicellular programs from large-scale spatial transcriptomics datasets. While existing approaches offer valuable tools, they are typically limited to single-sample analysis or rely on data types that lack crucial spatial proximity information (Tab. 1). For instance, methods like COMMOT^27^ and BATCOM^24^ model LRI within a single sample by incorporating competition or diffusion processes but do not identify overarching programs across multiple samples. Other approaches that do extract latent programs rely on different data sources, each with limitations. MOFAcell^37^ computes programs from bulk RNA-seq data and then maps them onto spatial samples, DIALOGUE^41^ matches LR pairs to programs post-extraction on the full transcriptome, and Tensor-cell2cell^38^ uses single-cell data, lacking the spatial dimensions. In contrast, our method uniquely integrates precise, spot-level interaction modeling, which includes neighborhood effects, diffusion, and competition, with the ability to extract robust, multi-sample programs. COMPOTES integrates in a hybrid fashion two common approaches for CCC analysis^27–29^. First, we modeled the LRI at the level of each spot, for each sample. (Fig. 1A). To achieve this, we took into account each ligand expression in neighboring spots by modeling a diffusion process depending on the type of ligand considered. Using CellChatDB^9^ annotations we either only consider the ligand expression within a spot (no diffusion), or add the weighted expression of its first or second layer of neighboring spots (if diffusing, see Methods). We then applied a competition step by normalizing the ligand score by the sum of all ligands scores that could bind to a receptor of interest. Finally, we computed the product of the obtained ligand score with the receptor expression. The second step of this method consists in a global extraction of latent multicellular programs (Fig. 1B). We applied a non-negative CANDECOMP /PARAFAC (CP) decomposition on a matrix containing every spot from multiple samples. This yielded programs describing the activation of a weighted list of ligand-receptor pairs in space. In particular, these programs can represent niche biological processes existing in very specific cellular contexts across multiple patients.

**Figure 1.**
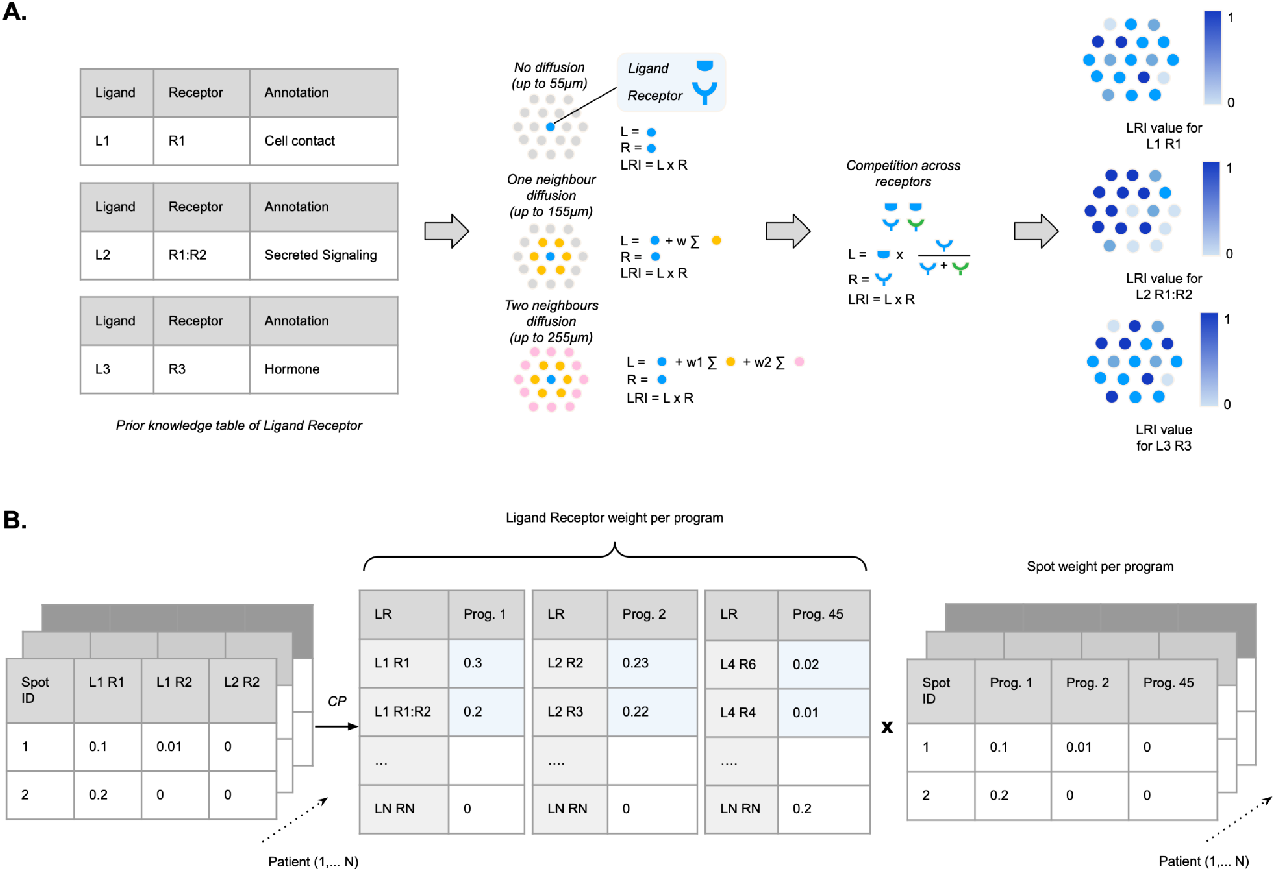
Method overview. **A.** Ligand Receptor Interactions (LRI) computing. From the prior knowledge table three types of LR interactions are defined: short, medium and long. The short ones do not diffuse (up to 55µm, intrinsic to the spot), the medium diffuse to the first layer of neighbours (up to 155µm), the long ones can diffuse to the second layer of neighbours and beyond (up to 255µm). A competition term is added, which penalises ligands associated with receptors known to bind multiple ligands expressed in the neighborhood. The contribution of each neighboring spot is weighted based on a circular diffusion mode (see Methods). Hence, each spot has a LRI value, as described by the right heatmap. **B**. From the LRI, K programs of communications are extracted using the CP decomposition. These programs are represented as a list of ligand receptor weights and as a list of spot weights.

To validate that COMPOTES accurately recovers expected signal, we simulated 10 Visium slides, populated with 11 hypothetical cell types (A, B… K) interacting with each other through LR pairs (Fig. 2A, see Methods). To demonstrate the advantage of using spatial data, we designed the simulation so that two of the six potential interacting cell type pairs were never in close spatial proximity, inducing potential false positives. COMPOTES correctly identified K=4 programs corresponding to the four true, spatially-proximal interactions, and correctly rejected the two non-interacting pairs (Fig. 2B). The method also successfully identified an interaction specific to a subset of samples. In contrast, when we ran tensor-cell2cell on the same data, it identified K=6 programs, incorrectly recovering the two pairs that lacked spatial proximity, generating false positives (Fig. 2C). We further tested this on a larger-scale simulation of 50 slides with greater cellular and LRI complexity. When running both methods for K=2 to K=40 programs, COMPOTES progressively recovered the true interactions, reaching the full set at K=40. Conversely, tensor-cell2cell recovered false positive signals at nearly the same rate as true positive signals (Fig. 2D). These simulated experiments suggest that COMPOTES effectively eliminates the false positives inherent to non-spatial methods without increasing false negatives, thereby accurately capturing true multicellular communication patterns.

**Figure 2.**
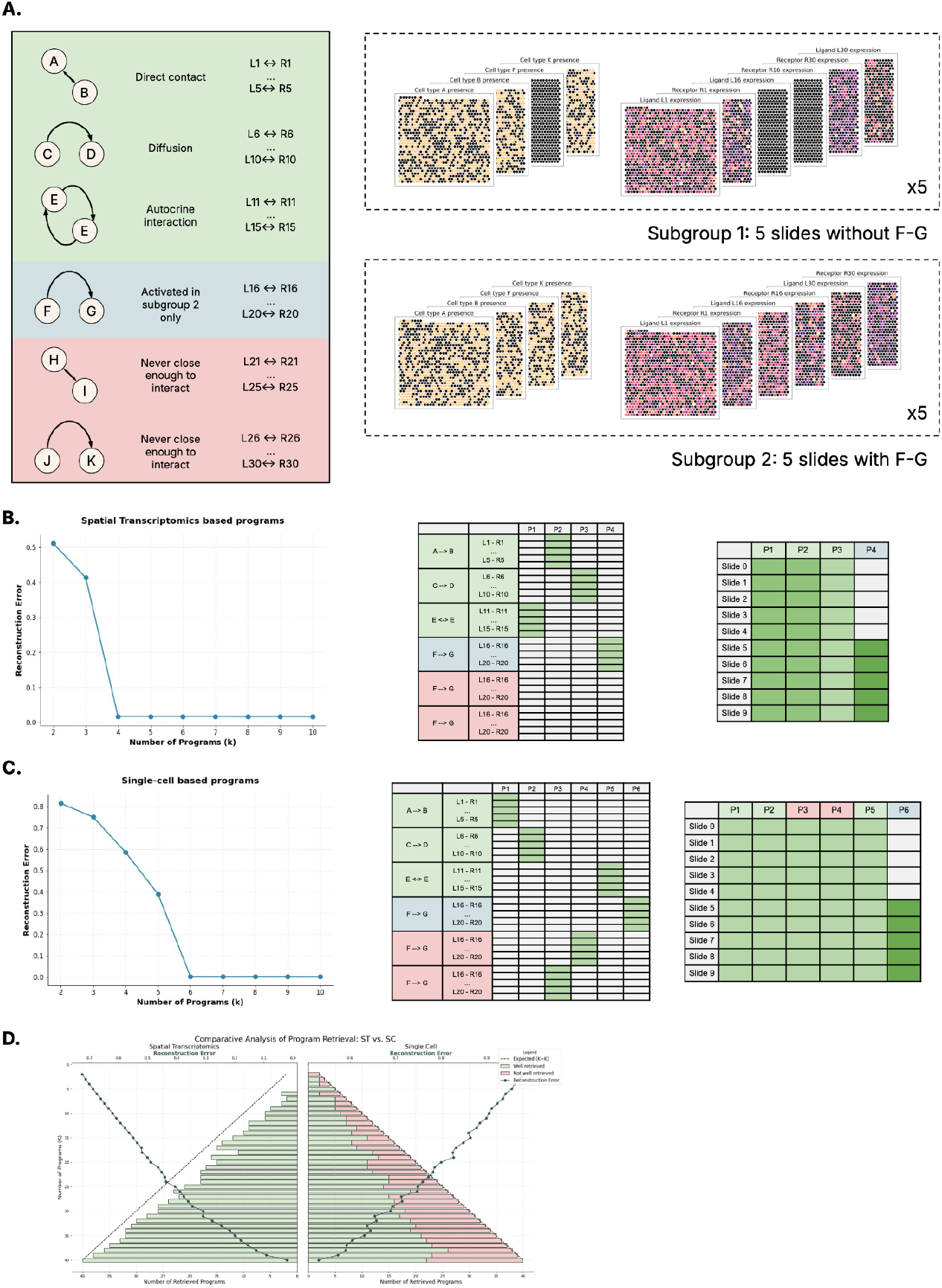
Validation of the method on synthetic data. **A.** Simulation of spatial transcriptomics slides. 10 slides are generated with 1000 spots arranged in an hexagonal grid. Each spot can contain multiple cells amongst 11 pre-defined cell types (A, B, … K). Those cell types can interact with each other such as described in the tabular (left) through 5 LR pairs for each interaction. Each cell type randomly expresses ligand and/or receptor following a Poisson law since sequencing read counts are discrete events that can be modeled as independent samples from a transcript pool. Depending on the interaction, ligands can diffuse to the immediate neighbors of each spot. If a ligand and the associated receiver cell types are present in the same spot, the receptor expression is increased by the expression of the ligand to model the LR interaction. Some cells are placed in the spots so that they are never close enough to interact with their associated interaction (in red on the left, HI and JK). Some cells are only placed in half the samples (in blue on the left, FG) so that the programs associated with these cell types should appear selectively among these samples. **B**. Results obtained when applying COMPOTES on the simulated dataset. On the left is the reconstruction error, showing a first minima for K=4. In the middle are the weights of the K=4 programs for the LR pairs of each possible cell type interaction. On the right are the weights of the K=4 programs for each of the 10 slides. **C**. Results obtained when applying tensor-cell2cell on the simulated dataset. To simulate single-cell data, each spot of the spatial transcriptomics slides are separated into individual cells, based on the cells they contain. On the left is the reconstruction error, showing a first minima for K=4. In the middle are the weights of the K=4 programs for the LR pairs of each possible cell type interaction. On the right are the weights of the K=4 programs for each of the 10 slides. **D**. Results obtained on a larger-scale simulated dataset. We compare the number of accurately retrieved programs (True Positives, in green) vs the number of programs that should not have been retrieved (False Negatives, in red) for both approaches.

**Table 1.** Comparison to existing methods.

| Method | Core Computational Engine | Primary Input Data | Multi-Sample Integration | Spatial Handling | Output Resolution |
| --- | --- | --- | --- | --- | --- |
| COMPOTES | Neighborhood LRI Score + CP Tensor Decomposition | ST (Visium) | Native (Matrix Factorization) | Neighborhood-based Score | Multi-sample factors; Spot-level LRI scores |
| Tensor-cell2cell | CP Tensor Decomposition | scRNA-seq | Native (Tensor Factorization) | Post-hoc (Niches as contexts) | Multi-sample factors |
| MOFAcell | Group Factor Analysis (MOFA) | scRNA-seq | Native (Matrix Factorization) | Post-hoc (Views can be spatial) | Multi-sample "multicellular programs" |
| DIALOGUE | Penalized Matrix Decomposition + Multilevel Models | scRNA-seq or ST | Native (Matrix Factorization) | Post-hoc (Niches as samples) | Multi-sample "multicellular programs" |
| COMMOT | Collective Optimal Transport | ST or spatially-annotated scRNA-seq | Single-Sample Focus | Global Optimal Transport | Spot/cell-level CCC matrix; Signaling vector fields |
| BATCOM | Bayesian/Frequentist Generalized Linear Model | ST | Single-Sample Focus | Explicit Distance Decay | Cell-type pair communication strength (with p-value) |
| HoloNet | Multi-view Graph Neural Network (GNN) | ST | Single-Sample Focus | Explicit Distance Decay | Functional LR pairs; Cell-type FCE network |
| DeepCOLOR | Variational Autoencoder (VAE) | ST + scRNA-seq | Single-Sample Focus | Recovers single-cell spatial map | Single-cell colocalization network |
| NICHES | LRI Product Matrix + Dimensionality Reduction | scRNA-seq or ST | Single-Sample Focus | Neighborhood Filtering | Single-cell to single-cell interaction matrix |
| scSeqComm | Permutation-based Scoring + Pathway Analysis | scRNA-seq | Single-Sample Focus | None (validated with ST) | Inter- and intra-cellular signaling scores |
| NATMI | Expression & Specificity-based Scoring | scRNA-seq or Bulk | Pairwise Comparison | None | Cell-type pair interaction scores |

### Application to a large multi-indication dataset

To evaluate our method, COMPOTES was applied to the MOSAIC dataset^42^, a large multi-indication and multimodal cancer dataset (see Methods). Specific sets of programs were computed for each of the 7 MOSAIC indications individually, and for all aggregated indications. This translates to 3.5 million spots across 1084 samples from 4 centres including: 248 non-small cell lung cancer (NSCLC) samples, 240 ovarian cancer (OV) samples, 154 diffuse large B cell lymphoma (DLBCL) samples, 146 muscle-invasive bladder cancer (MIBC) samples, 118 breast cancer (BC) samples, 112 glioblastoma (GBM) samples and 66 pleural malignant mesothelioma (MESO) samples.

First, we analyzed the programs obtained with each single indication. Using 146 baseline samples from the MIBC cohort, we generated sets of programs for different numbers K of programs. We selected K=45 programs using relevant heuristics, including reconstruction error, cell type involvement, ligand-receptor (LR) pair diversity, and program specificity (see methods for more details). These programs were then grouped into 3 categories based on the cell types communicating and the types of ligand-receptor pairs mediating the interaction: (1) First, the Immune-related programs group was defined by programs involving a high proportion of immune cell types as sender and/or receiver cell type; (2) Second, ECM mediated group was defined by programs characterized by LRI annotated as ECM-receptors; (3) Finally, Growth factors group, proliferation and stromal-malignant interactions group regrouped the residual programs after inspection of the LRI and cell types characteristic of these factors (Fig. 3A, Methods, Supp. Tab. S1-2). Importantly, clustering of both programs and samples revealed no particular enrichment in any of the three MIBC cohorts from different centers, confirming a shared biological signal (Fig. 3B).

**Figure 3.**
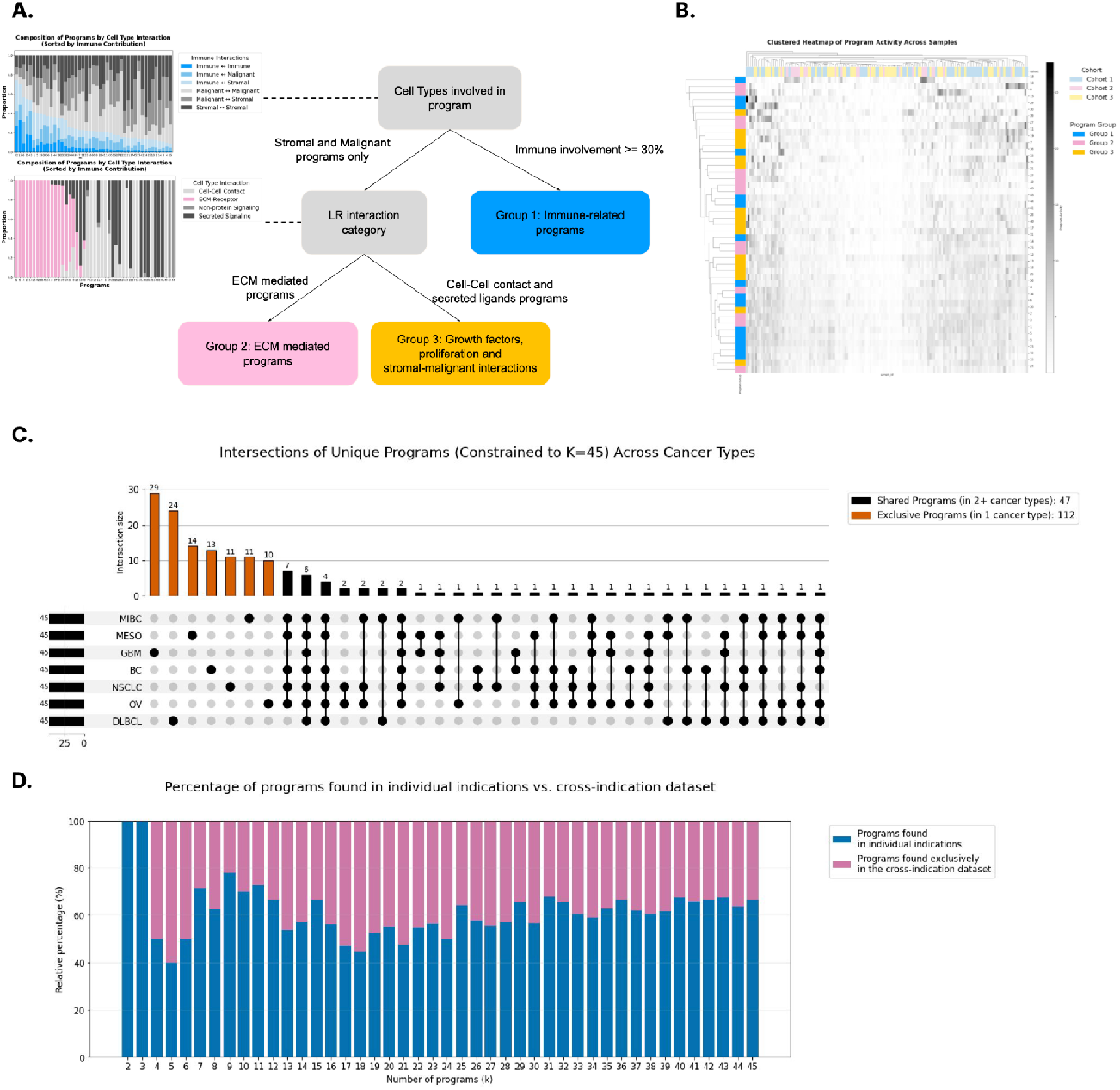
Results Overview. **A.** Analysis of the set of K=45 programs obtained with the MIBC cohort. For each program, we compute the associated cell types and the types of LR pairs mediating the interaction (see methods). Using those attributes, we define 3 comprehensive groups of programs. **B**. Clustering of the MIBC programs across the input samples annotating the center from which samples were harvested and the 3 groups of programs described in panel A. **C**. Analysis of the overlap of the obtained factors computed with each single cohort. The upset plot shows the intersection of the programs matched through cosine similarity of their respective weighted LR pairs (see methods) between indications. **D**. Analysis of the similarity of the programs obtained when grouping the cohorts together compared to using the cohorts separately. The barplot shows the percentage of programs that were found in individual indications (blue) and the percentage of programs that were not found (pink) across different values of K=2 to K=45.

Second, we compared the results obtained for each single indication, again selecting sets of K=45 programs, using cosine similarity on the vector of LR-pairs weights to match (if cosine similarity above 0.8) programs between experiments. We observed that each indication shared numerous programs, while also yielding indication-specific ones (Fig. 3C, Supp. Tab. S2). NSCLC, MIBC, MESO, BC and OV yielded the highest number of similar programs with 32-35 similar programs found in at least another indication and 19 programs found in all these 5 indications. By contrast, GBM and DLBCL yielded the most singular programs, with only 16 to 21 similar programs. This is consistent with the fact that GBM and DLBCL have a unique biology both in terms of cell of origin but also TME contexts as compared with non-CNS (non-Central Nervous System) solid tumors. Therefore these indications seem to be associated with specific CCC patterns not shared with other indications.

Last, we investigated the difference in the obtained results when computing programs across every indication concatenated. We observed that while most programs matched the ones found across the 7 indication specific experiments (∼60%), a significant fraction was new (Fig. 3D). This can be explained by the increased complexity of existing biological niches in the entire dataset, needing a higher K to decompose biological programs at the same granularity level.

Overall, we note that the overlapping programs found across different indications were mostly characterized by LR pairs with the same receptor (such as CD44, SDC1, SDC4 or DAG1). However, some programs with complex LRI composition identified in the MIBC analysis, such as program 15, were also retrieved in other indications. (Supp. Tab. S2). Lastly, some programs were entirely indication-specific, like the MIBC’s specific program 7. Thus, we note that this method is able to yield both pan-cancer general programs that characterize common phenomena across different disease contexts; and indication specific-programs that characterize cellular communications disease specific.

### Program 15 identifies a mounted yet inefficient anti-tumoral immune response

We used our method to analyze Program 15 (P15), a robust communication network found in six of the seven cancer types in our study (Supp. Tab. S2). The program’s stability was further confirmed in MIBC experiments, where it consistently appeared across multiple K (Supp. Tab. S3). Several statistical associations between P15 activity and patient level or spot level information (Supp. Tab. S2 and S4, respectively) were further computed to highlight COMPOTES capabilities dissecting complex cancer biology.

This program represents a recurrent ecosystem of interacting cells and is defined by the coordinated activity of numerous LR pairs, central to immune function (Supp. Fig. S3A). The involved LR pairs encompass a wide range of processes, including innate-adaptive crosstalk (IL16-CD4)^43^, leukocyte migration (CCL5-CCR5)^44^, immune checkpoint signaling (NECTIN2-TIGIT)^45^, and the clearance of apoptotic cells (C3-ITGAX:ITGB2)^46^. This diversity marks P15 as a key hub of immune activity.

To dissect the cellular architecture of this network, we integrated matched single-cell data. This revealed a complex interplay between fibroblasts, T/NK cells, and monocyte-macrophages (MoMacs), with a clear pattern: ligands were expressed by various cell types, but receptors were predominantly found expressed in MoMacs, positioning them as the primary signal receivers (Supp. Fig. S3B-C). Spatial analysis confirmed these findings, showing the strongest co-localization between P15 activity and MoMacs when compared to other programs, followed by T/NK cells and DC (cosine similarities=0.36, 0.20 and 0.15, respectively; Supp. Fig. S4A, Supp. Tab. S4). P15 activity was significantly associated with the MIBC Basal patient subtype (Supp. Fig. S3D), using the sum of spot activity at the patient level. In addition, we also found P15 to be spatially concentrated at the tumor-stroma interface (Supp. Fig. S3E) and was significantly higher in the vicinity of T-cell aggregates (Supp. Fig. S3F-G) inferred from histopathological images using an in-house model (see methods). This suggests a relationship between P15 and immunologically “hot” regions, knowing that the Basal tumors in MIBC are more immune-infiltrated than Luminal tumors^47^. In line with Basal MIBC patients responding better than Luminal patients to immune checkpoint inhibitors (ICI)^48^ which re-invigorate a stalled immune response in tumours, we observed that P15 was significantly associated with the *Immune Response to IO T-cell inflamed* pathway previously published by Cristrescu *et a*l^49^ (cosine similarity = 0.15, Supp. Fig. S3B). A representative patient sample visually confirmed that P15 activity peaks in immune-rich stromal regions adjacent to the tumor core, where T-cell aggregates are detected (Supp. Fig. S3G).

Downstream functional analysis revealed the paradoxical dual nature of P15. On one hand, its activity strongly correlated with canonical pro-inflammatory pathways, including JAK-STAT, TNFɑ, and NF-κB, and an “IFN-ɣ response” gene signature (cosine similarity = 0.40; Supp. Fig. S4B-C). This indicates the presence of a robust, active anti-tumor immune response. On the other hand, the program was as strongly associated with signatures of immune dysfunction. In this regard, we observed high similarity to signatures of T-cell exhaustion^50^ (cosine = 0.31) and immunosuppressive tumor-associated macrophages (TAMs)^51^ (cosine = 0.28, highest program associated) (Supp. Fig. S4B). This suppressive environment was further evidenced by an association with regulatory T-cells (Tregs) and type-2 conventional dendritic cells (cDC2s) (Supp. Fig. S4D), an axis known to blunt effective anti-tumor immunity^52^.

In summary, P15 captures the signature of a stalled, exhausted anti-tumor immune response. It identifies a spatially organized TME where the immune system has been successfully mobilized (as highlighted by an active pro-inflammatory signaling and proximity to T-cell aggregates) but is simultaneously being held in check by powerful suppressive mechanisms. The network highlights MoMacs as central players, simultaneously participating in the anti-tumor response while being associated with a pro-tumoral, immunosuppressive phenotype. This finding pinpoints a critical state of immune equilibrium where activation is actively counteracted, preventing effective tumor clearance.

### Program 7 defines a *KMT2D*-linked temporarily and spatially confined MIBC-specific proliferation engine

We next sought to better investigate the nature of indication-specific programs identified by COMPOTES, as they potentially identify unique cellular communications networks. For this, we focused on Program 7 (P7), a program found only in MIBC tumors. This Program displays consistent stability across multiple factorization experiments (Supp. Tab. S2-S3) and is part of Group 3, involving growth factors, proliferation, and stromal-malignant interactions (Supp. Tab. S2). P7 results from several LR pairs (Fig. 4A) operating along four main axes: (1) a ‘neuro-developmental axis’ that remodels the TME^53^ through axon guidance pathways (Semaphorin-Plexin^54^, Ephrin-Eph^55^, Netrin-UNC5^56^), promotes angiogenesis, and enforces immune exclusion^57^; (2) a ‘stromal and adhesion axis’ driven by non canonical WNT5A^58^ and adhesion molecules (LRFN4 or Nectins^59^) that activates cancer-associated fibroblasts (CAFs) and stabilizes intercellular contacts^60^ fostering a fibrotic, therapy-resistant niche^61^; (3) an ‘immunomodulatory axis’ (TNFSF15/TNFRSF25) that fine-tunes lymphocyte homeostasis and function^62^; (4) a ‘metabolic-immune axis’ (DHCR8/LXR), fueling cell proliferation while potentially creating an immunosuppressive and anti-inflammatory, cholesterol-rich TME^63^.

**Figure 4.**
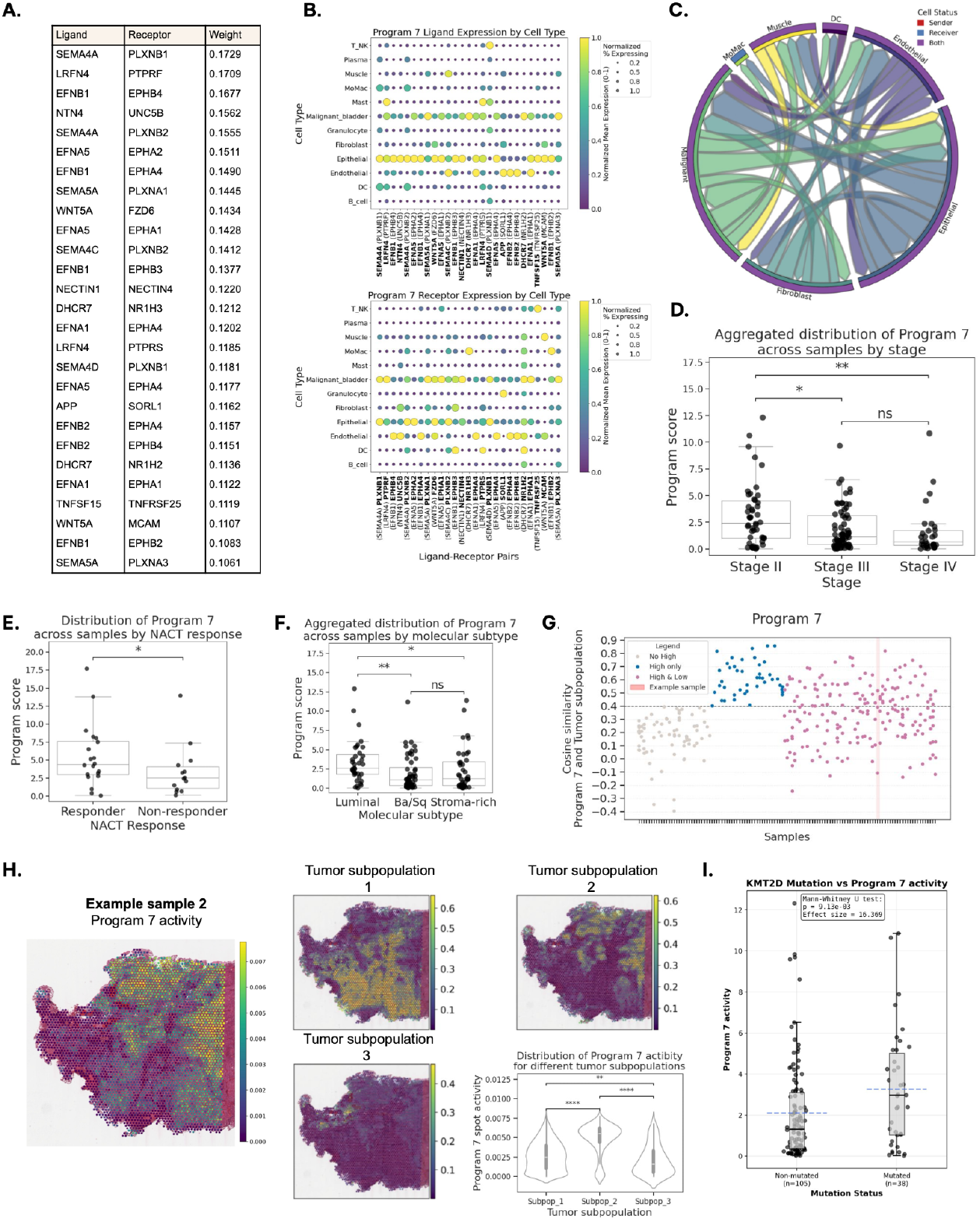
Analysis of the Program 7 (P7). **A.** Top LR pairs of P7 (weight ≥ 0.1). **B**. For each of the top LR pairs, mean expression and percentage of cell types expressing each ligand (top) and receptor (bottom) in single-cell data. **C**. Cellular network of P7. **D**. Association of P7 with the stage T at the sample level. E. Association of P7 and the response to NACT treatment. **F**. Association between P7 expression and the deconvolution fractions of malignant subpopulations for each sample. **G**. Example of one of the samples. The following are shown across spots: expression of P15; normalized deconvolution fractions of the 3 observed tumor subpopulations; violin plot showing the activity distribution of P7 in the three subpopulations. **H**. For the example sample, distribution of P7 expression across spots of each of the 3 tumor subpopulations. **I**. Boxplot indicating the activity of the P7 in each sample, either Non-mutated (left) and Mutated (right) for *KMT2D*. p-value was calculated with a one-sided Mann-Whitney statistical test.

The signaling network has a highly connected group of cell types, with stromal and cancer cells predominantly expressing the ligands and receptors of P7 (Fig. 4B). Sender-receiver modeling revealed a network driven by both paracrine (Epithelial ↔ Malignant) and autocrine (Malignant → Malignant, Epithelial → Epithelial) signaling loops, forming a self-reinforcing communication engine. (Fig. 4C). Endothelial and fibroblast cells also act as significant receivers. Spatial transcriptomics analysis confirmed the co-localization of these cells, showing a strong association between P7 activity and malignant and epithelial cells. (average cosine similarity of 0.46 and 0.37, respectively) (Supp. Fig. S5A).

To understand P7 effects, we analyzed its spatial association with cancer pathways and clinical data. P7 activity correlated strongly with multiple classic cancer hallmark signatures (PROGENy), including PI3K, VEGF, MAPK, Androgen, Estrogen, and WNT pathways (average cosine = 0.56, 0.57, 0.53, 0.48, 0.54, 0.49 respectively) (Supp. Fig. S5B, Supp. Tab S4). This convergence of multiple oncogenic forces, whereby hormone signaling activates core growth pathways (PI3K and MAPK) and in turn promotes angiogenesis (VEGF), suggests a robust network that synergistically enhances proliferation (MAPK, PI3K)^64^ and migration (VEGF, WNT)^65 66^. This is further supported by a similar enrichment in manually curated gene signatures from literature related to fundamental cancer hallmarks such as RTK^67^, Hippo^67^, NOTCH^67^, PI3K signaling^67^, dysregulation of the TP53 pathway^68^, and cell cycle^68^ (average cosine = 0.31, 0.32, 0.34, 0.35, 0.39, 0.40 respectively). In particular, cell proliferation resulted to be strongly associated with P7: out of the three custom signatures extracted from literature related to cell cycle used in this analysis (Cell cycle Proliferation rate Bagaev et al.^51^, Cell cycle Sanchez-Vega et al.^67^, Cell cycle Telomere Maintenance and Cellular Process Cell Cycle Blum et al.^68^), P7 was the program with the strongest association across all 45 programs identified by COMPOTES in MIBC (Supp. Fig. S5C, Supp. Tab S4).

When examining the patient-level activity of P7, a significant increase was observed in patients at the T2 stage, suggesting the network’s role in the early invasive steps of MIBC. (Fig. 4D, Supp. Tab S2). P7 also tends to be more activated in patients who had a positive response to neoadjuvant chemotherapy (NACT) (p.val = 0.03; Fig. 4E, Supp. Tab S2), which is supported by the program’s associations with several chemotherapy response signatures (Kato et al ^69^, Takata et al ^70^ and Yu et al ^71^ signatures; cosine similarity of 0.23, 0.26 and 0.27 respectively) (Supp. Fig. S5E, Supp. Tab/S3). This is consistent with the pathway analysis, as highly proliferating cancer cells are more likely to be in vulnerable phases of the cell cycle and thus more susceptible to the damage induced by chemotherapy drugs. Additionally, P7 is significantly associated with Luminal MIBC subtypes, which are also known to be more responsive to NACT. (Fig. 4F, Supp. Tab. S2).

Through the integration of spatial and single-cell data, we identified transcriptionally and spatially distinct cancer cell populations, which can be individually correlated with the activity of programs identified by COMPOTES (see methods). This revealed that P7 had the strongest intra-tumoral heterogeneity of activation among all the programs identified in MIBC, with 51% of patients displaying both activating and non-activating subpopulations (Fig. 4G-H, Supp. Tab S2). Similarly to P15, we also explored the spatial localisation of P7 activity with regard to tumor islet core, tumor margin and stroma and found that this program tends to be activated at the core of the tumor, which can also be reconciled with pathway analysis, and especially angiogenesis pathways being enriched via the semaphorin LR pairs (Supp. Fig. S6A; Fig. 4H).

We then explored the association between the patient level activity of P7 and oncogenic driver gene alterations, including both somatic mutations or copy-number alteration. This allowed us to identify that *KMT2D* mutations are significantly associated with patient level activity of P7 (p-value = 0.0091; q-value = 0.047; Fig. 4I, Supp. Fig S6B).

We finally explored the reason for the representation of epithelial cells as a major component of the P7 sender–receiver network (Fig. 4C). We considered the possibility that our annotation and deconvolution pipeline might classify as epithelial cells, cells in a juxtatumor tissue context being in a pre-tumoral dysplastic state, consistent with the well known existence of cancer driver gene mutations in normal-appearing urothelium, of which the most frequently mutated included chromatin remodeling genes, such as *KMT2D*^72^. To investigate this, we explored a specific example of a sample (Supp. Fig S6C) with a clear area of epithelial cells identified through deconvolution characterized by a cnv_score similar to regions with high malignant cell deconvolution fraction. Interestingly this epithelial rich region had a higher P7 activity than the malignant region. Histopathology investigation led to the identification of an epithelium with atypia and architectural disorder without loss of maturation and polarity, suggesting dysplastic cells that could already be characterized by malignant features. To validate this hypothesis, we assessed the relationship between chromosomal instability (cnv_score) and P7 activity across tumor regions using linear mixed-effects models with random slopes and intercepts grouped by patient (Supp. Fig S6D). Cnv_score emerged as the dominant predictor of P7 activity (standardized β = 0.43, z = 14.50, p < 1×10^-47^), with an effect 7.1-fold stronger than tumor fraction (standardized β = 0.060, z = 2.71, p = 0.007), indicating that chromosomal instability is the primary driver of P7 activation. When we introduced an interaction term between tumor fraction and CNV score, we found a highly significant positive interaction (standardized β = 0.015, z = 11.90, p < 1×10^-32^), indicating that CNV effects on P7 activity are amplified in regions with higher tumor content. In this interaction model, the main effect of CNV score remained strong (β = 0.42, z = 14.27, p <1×10^-45^), while tumor fraction showed a moderate but significant direct contribution (β = 0.066, z = 2.97, p = 0.003). This supports the interpretation that P7 captures a tumor-intrinsic communication program tightly linked to genomic instability, and further raises the possibility that some epithelial annotations in high-P7 regions may reflect early tumoral, dysplastic epithelial states rather than normal epithelial cells.

In summary, our results delineate P7 as a unique and multi-faceted biological state specific to luminal MIBC. Its defining characteristic is a specific set of LR interactions, which we propose drive the activation of key cancer hallmarks. While similar hallmark activation may occur in other cancers through distinct pan-cancer or indication-specific programs, our data specifically underscore the unique set of LR pairs underpinning P7’s effects in luminal MIBC. P7 integrates neuro-developmental, stromal, immunomodulatory, and metabolic signaling pathways, collectively promoting a strong proliferative phenotype. Paradoxically, despite its association with numerous oncogenic pathways, this program is most active in early-stage tumors, is spatially restricted to the tumor core, and predicts a favorable response to NACT. Furthermore, P7 is significantly linked to inactivating mutations in the epigenetic modifier *KMT2D*. This association suggests that alterations in the tumor’s epigenetic landscape, specifically in the context of P7, would influence therapeutic response.

## Discussion

COMPOTES is a new hybrid method that combines single-cell and spatial-transcriptomics to analyze CCC. It uses matrix factorization on spatially colocalized ligand and receptor interaction to extract multicellular communication programs across entire cohorts of samples. This bridges the gap between accurate local CCC, and extraction of trends across multiple samples.

When applied to the MOSAIC dataset, our approach identified a collection of different communication patterns, some shared across different tumors and others unique to a specific disease. P15 represents a good example of a recurrent communication network across indications, with a dual character highlighting the balance of immune activation and suppression dictating tumor fate. The simultaneous presence of strong IFN-ɣ signaling, markers of T-cell and macrophage exhaustion suggests a state of adaptive immune resistance, where suppressive feedback loops, potentially driven by IFN-ɣ itself, neutralize the anti-tumor response. This underscores that simply boosting immunotherapy may be insufficient and that specific interactions within P15, like the NECTIN2-TIGIT and cDC2-Treg axes, are potential targets for new combination therapies.

Conversely, COMPOTES identifies a MIBC specific program P7 which paradoxically is rich in oncogenic pathways but strongly linked to favorable clinical outcomes, namely T-stage and NACT response. Therefore, we propose a “high-proliferation, high-vulnerability” model. The program’s proliferative activity is driven by multiple signals, particularly the Semaphorin-Plexin system. Semaphorin 4D (Sema4D), upregulated in bladder cancer, directly increases proliferation and motility^73^. This is amplified by synergistic signaling from Nectin-4 (via PI3K/AKT ^73,74^) and LRFN4 (via Ras^75^). This multi-pronged activation of growth pathways explains why rapidly dividing cells are more susceptible to cytotoxic chemotherapy, leading to a strong NACT response^69,73,75^.

Concurrently, growth is contained by intrinsic molecular brakes, like Netrin-1/UNC5B neuro-developmental pathway, functioning as an apoptotic “kill switch”. The P7 program’s highly proliferative cells are dependent on the Netrin-1 ligand for survival: in the presence of Netrin-1, the UNC5B receptor transduces a pro-survival signal, suppressing apoptosis.^76^ This dependency forces cells to stay in a Netrin-1-rich microenvironment to escape programmed cell death. This mechanism elegantly explains three of the program’s most striking features: its prevention of local invasion (enrichment in T2 stage), its confinement to the tumor core, and its high degree of intra-tumoral heterogeneity.

The program highlights an immune-cold niche, which is consistent with the Luminal molecular subtype. This is primarily driven by the metabolic-immune axis: upregulated cholesterol production fuels tumor growth while also creating a lipid-rich, immunosuppressive microenvironment. This effect makes the tumor non-inflamed, allowing it to evade immune surveillance^77,78^. The strong association with KMT2D mutations suggests a master regulatory event. *KMT2D* is a lysine methyltransferase and core component of the KMT2C/D-KDM6A complex, mediating the methylation of histone H3K4, generally associated with active enhancers and transcriptional activation and playing a pivotal role in the regulation of cell-lineage specific programs^79,80^. *KMT2D* is frequently mutated in cancers, including urothelial carcinoma, harboring loss-of-function mutations in key epigenetic modifiers^81,82^. While not enough to cause cancer on its own, its loss ‘primes’ the urothelium for transformation resulting in a global downregulation of the urothelial differentiation program, and augmented stem/progenitor potential^83^. P7 association with the Urothelium cell type (Supp. Tab. 4), suggests a ‘primed’ niche in MIBC patients.

The loss of functional KMT2D likely causes a widespread H3K4me-mediated dysregulation of the cellular transcriptome and activates a dormant developmental program in adult urothelial cells, providing a plausible mechanism for the multi-axial activation of Program 7. Furthermore, KMT2D is known to play a role in regulating neuro-developmental pathways^84^ including, plausibly, the axon guidance systems (Semaphorin, Ephrin, Netrin) that are central to Program 7. We propose that the somatic loss-of-function of KMT2D leads to the aberrant re-wiring of this entire dormant program, inducing a cell reprogramming which elegantly accounts for the simultaneous emergence of its disparate features as a coordinated unit linked to a specific genetic background. Additionally, KMT2D loss may also explain the increased drug sensitivity of P7 cells^85–88^. This could be due to increased epigenetic vulnerabilities, where the *KMT2D*-dependent altered chromatin state makes cells more susceptible to DNA damage that could occur via the disruption of DNA repair mechanisms^89^, making them more vulnerable to chemotherapy drugs. Yet, the exact mechanism would require further investigation.

In conclusion, Program 7 defines a distinct biological entity in early-stage, KMT2D-mutant, Luminal MIBC. Its favorable prognosis is likely due to its inherent vulnerabilities, namely extreme chemosensitivity and architectural limits, which outweigh its oncogenic potential. This suggests that tumors may need to downregulate this program to escape its fragility, making P7 a promising target for understanding NACT response and developing new therapies.

The non-exhaustive exploration of these two programs is representative of COMPOTES potential to decipher the cellular ecosystems driving tumor progression. However several avenues of improvement to our approach can be highlighted. While our current method allows us to analyze the connection between CCC programs and cellular functions after computation, other recent methods like HoloNet^11^ or scSeqComm^90^ directly incorporate this mechanistic association into their model. More generally, this integration could extend to other types of information beyond cellular functions, such as disease states. For instance, Tensor-cell2cell natively integrates the latter using their multi-dimensional tensor. While this is possible with the proposed methods, it faces computational scalability issues: LRI calculation must be performed for every spot in every sample, which leads to the decomposition of a large matrix. With 146 samples and 500 000 spots, we are already reaching the memory limits of a standard 16GO VRAM GPU. As such, scaling to either more samples or adding other dimensions to include sample-level or spot-level annotations would correspond to a computational bottleneck. Finally, our current analysis and results are constrained to the spot-level resolution offered by this 10x Visium technology but our method could theoretically be extended to platforms with higher levels of resolution, such as 10X Visium HD or FISH-based methods.

## Methods

### Data

#### MOSAIC

MOSAIC is an ongoing initiative led by Owkin and 5 hospitals aiming to generate the world’s largest spatial atlas in cancer. It collects and generates six data modalities (extensive clinical data, H&E slides, 10X Visium spatial transcriptomics, single-nuclei transcriptomics, bulk RNAseq, and whole-exome sequencing). The technology used is Visium V2 Cytassist, applied to FFPE samples, and the captured area covers 6.5 mm x 6.5 mm.

Here we leveraged samples originating from 7 indications and 4 centres, distributed across indications as follows: n=248 lung samples, n=240 ovarian samples, n=154 Diffuse large B cell lymphoma (DLBCL) samples, n=146 Muscle-invasive bladder cancer (MIBC) samples, n=118 breast samples, n=112 glioblastoma (GBM) samples and n=66 mesothelioma samples.

In particular, we leveraged n=146 samples from the MIBC dataset. The samples are originating from 3 of the founding hospitals in the MOSAIC consortium: Lausanne University Hospital - CHUV (64 samples), Charité - Universitätsmedizin Berlin (25 samples), Universitätsklinikum Erlangen (57 samples). The detailed methods associated with the MOSAIC data are available in the associated paper^42^. Data are part of the MOSAIC consortium dataset, a non-interventional clinical trial registered under NCT06625203.

#### CellChatDB

CellChatDB is a database including literature-supported ligand-receptor interactions in both mouse and human species. The majority of ligand-receptor interactions were manually curated using the KEGG signaling pathway database^91^.

#### Simulated data

To validate our method, we generate pseudo-Visium slides and pseudo-single-cell data. For each slide, N spots are first arranged in a hexagonal grid. Then, a list of cell types (CT_1_, … CT_M_) is defined. Each cell type has an associated probability p_CTi_ to be present in a spot. For each spot, the presence or the absence of each cell type is computed. Then pairs of interacting cell types (CT_i_ <-> CT_j_) are defined, with associated LR pairs mediating those interactions. The ligands can mediate short interaction within the same spot or diffuse to the neighboring spots. The baseline expression of ligand and receptor is modelled using a Poisson distribution with parameters λ_ligand_ and λ_receptor_. For each spot, for each cell type emitting a ligand or having a receptor, a baseline expression is computed for every ligand and receptor. Then, for diffusing ligands, the diffusion of the ligand is modelled and the diffused value (initial value divided by a fixed number) is added to the neighboring spots. Lastly, the interaction between ligand and receiver cells is modelled in the following fashion: if in a given spot, a receiver cell and the associated ligand are present at the same time, the expression of the ligand is added to the expression of the associated receptor.

With this framework, we model the two following cases:

1. Simple case. 11 cell types (A, B, …, K) interacting in the following fashion:
  - A→B, 5 LR pairs with diffusion
  - C→D, 5 LR pairs without diffusion
  - E↔E, 5 LR pairs without diffusion
  - F→G, 5 LR pairs with diffusion, but those cell types are only present in half the samples
  - H→I, 5 LR pairs with diffusion, but those cells are never placed close enough to interact
  - J→K, 5 LR pairs without diffusion, but those cells are never placed close enough to interact We model 10 slides of 1000 spots with parameters lambda_ligand=20 and lambda_receptor=5
2. Case with many interactions. 275 cell types interacting the following fashion:
  - 10 pairs of cells interacting with 5 diffusing LR pairs
  - 10 pairs of cells interacting with 5 not diffusing LR pairs
  - 10 pairs of cells with autocrine interacting with 5 LR pairs
  - 10 pairs of cells interacting with 5 diffusing LR pairs in only half the sample
  - 10 pairs of cells interacting with 5 diffusing LR pairs but those cells are never placed close enough to interact
  - 10 pairs of cells interacting with 5 diffusing LR pairs but those cells are never placed close enough to interact

We model 50 slides of 1000 spots with parameters λ_ligand_=20 and λ_receptor_=5

### LRI computation

The interaction between ligand and receptor is defined as the product of their normalized expression. The product is a common communication score, as it outputs a continuous value and will be set to zero if one of the measured transcripts is missing.

In the context of spatial transcriptomics, we consider that some ligands are membrane-bound and thus will only be interacting inside their spot; while other ligands are soluble molecules, able to interact within their spot and to travel a given distance away from it. In this second category, we further separate ligands able to travel a medium distance, which we model as able to travel from their spot to the immediate neighboring spots (up to ∼200um); from ligands able to travel further, which we model as able to travel from their spots to the second layer of neighboring spots (up to ∼400um). We did not take into account ligands that travel in the bloodstream and reach distant receptors.

The total amount of ligands that can reach the spot itself is determined by the physical phenomenon of diffusion. The contribution of each neighboring spot is weighted based on a circular diffusion model, where: 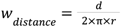. d represents the spot diameter (55μm), r is the distance from neighbor to target spot, and π is the mathematical constant pi. This model assumes that ligands emitted by a neighbor diffuse in a circular manner, with only a fraction reaching the target spot based on the distance and on the spot’s radius. The spatial layers consider immediate neighbors (6 spots at 100μm) and second-layer neighbors (12 spots at 100√3μm and 200μm). The methodology assumes uniform spot sizes (55μm diameter), standard Visium array geometry for neighbor distances, and isotropic diffusion in 2D space. The maximum interaction distance is capped at 2 layers (18 neighbors).

Thus we define the expression of the ligands as:

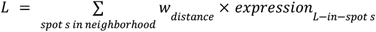

The receptors are membrane proteins attached to the surface of the cells. Thus, the expression of the receptor is taken as the expression in the considered spot:

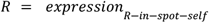

Therefore, the LRI strength is considered as the product of the expression of the ligands in a neighborhood multiplied by the expression of the receptors in the spot itself. *LRI* = *L* × *R*. The ligand and the receptor can form multimeric complexes. In this case, we hypothesize that the expression of the complex is equal to the minimum expression of all its composing elements. This ensures that all components are present for the interaction to occur.

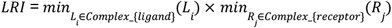

### CCC computation

We start from a pre-computed LRI matrix X spanning all N_s_ spots across the considered collection of samples and all N_LRI_ interactions considered. To extract multicellular groups of LRIs, we apply the non-negative CANDECOMP /PARAFAC (CP) decomposition^92–94^. The CP decomposition is a type of tensor rank decomposition, in which a given input tensor X of an arbitrary dimension n, is approximated as a sum of k tensors of rank 1 as follows:

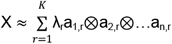

In the specific case of n = 2 dimensions, as is our case, it can be noted that this decomposition has a similar output to a non-negative matrix factorisation (NMF):

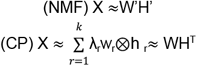

In practice, we use the implementation of the TensorLy python library which uses a Hierarchical Alternating Least Squares update algorithm^95^. This implementation minimizes the following problem:

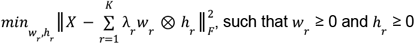

Where ║.║ _F_^2^ is the Frobenius norm.

It is to be noted that the final outputs, referred to as programs, are normalized individually using the L2 norm.

### Cell type assignment

The method utilizes single-nuclei RNA sequencing (snRNAseq) data to estimate the mean expression profiles of cell types, and assign weights to sender and receiver cells from program’s L/R loading.

First sncRNAseq count data were processed into a pseudobulk expression matrix. Then the different LR pairs were used to calculate the different cell type communication potential. For every selected L/R complex:

- The aggregate expression level of the ligand gene(s) was computed for each cell type. This yields a vector representing the potential “sending” capacity of each cell type for that ligand set.
- Similarly, the aggregate expression level of the receptor gene(s) was computed for each cell type, yielding a vector representing the potential “receiving” capacity.
- If the sum of expression across cell types was greater than zero, both the ligand and receptor expression vectors are independently normalized to sum to 1. This normalization distributes the total sending or receiving potential proportionally across the contributing cell types.

Therefore, for a given L/R complex, the potential communication score between a sender cell type and a receiver cell type was calculated as the product of the normalized ligand expression in the sender cell and the normalized receptor expression in the receiver cell.

This interaction score was then weighted by multiplying it by the absolute value of the loading of that specific L/R complex onto the program currently being analyzed. This weighting ensures that L/R interactions more strongly associated with the program contribute more significantly to the program-specific communication network.

Finally, the weighted communication scores for all L/R complexes associated with a given program were summed for each possible sender-receiver cell type pair (S,R). If the total sum of the aggregated matrix was non-zero, the entire matrix was normalized to sum to 1, providing a relative map of predicted communication flow between cell types specific to that program.

### Ablation study, benchmarking and method optimisation

To validate our approach and determine an optimal number of programs (K), we performed a comprehensive ablation and benchmark study. We compared our full model against two ablations, without competition or without diffusion, a negative control using randomly permuted gene expression, and a single-cell method (Tensor-cell2cell). Using 146 baseline samples from the MIBC cohort as input, we computed programs for K ranging from 2 to 100 and evaluated them on four key metrics: reconstruction error, cell type involvement, ligand-receptor (LR) pair diversity, and program specificity.

We observed that the reconstruction error steadily declines as K increases across all settings (Supp. Fig. S1A). The sharper initial drop for the single-cell approach can be attributed to its less sparse input data, a consequence of the averaging by cell types done in its processing. However, this approach also led to an under-representation of stromal cells in retrieved programs, highlighting the superiority of spot-level data in capturing the complete cellular ecosystem (Supp. Fig. S1B). Furthermore, while our full model and its ablations showed a balanced cell type inclusion, the ablation settings introduced biases in the representation of communication modes (Secreted Signaling, ECM-Receptor, Cell-Cell Contact). For instance, removing the competition component led to an over-representation of ECM-receptor interactions (Supp. Fig. S1C-D).

To assess program specificity and potential overfitting, we analyzed the programs’ complexity (Supp. Fig. S1E). An abundance of simple, single LR pair programs, could indicate overfitting. In our primary model, these simple programs began to appear around K≈40. This threshold appeared much earlier in our random permutation negative control, suggesting it overfits by smoothing signals specific to cellular niches. Indeed, the few programs found in both the main and permuted settings corresponded to broadly expressed genes like glycoproteins and collagens. The single cell approach identified a higher proportion of programs with multiple relevant LR pairs. However, as mentioned in the introduction, this method averages the signal of LR pairs per cell type, consequently reducing the resolution of particular niches that could be characterized by a very limited set of LR pairs representing its cellular interactions. Hence, in this context, this doesn’t represent a higher factorisation capability but a complex mixture of signals, resulting in the lack of resolution to identify particular cellular niches. These observations informed our selection of an optimal K as a trade-off: K<10 provides a rapid decrease in reconstruction error; K between 10 and 45 adds diverse and complex programs; while K>45 introduces signs of overfitting and incurs significant computational costs. Therefore, we chose K=45 as it optimally balances model performance, program interpretability, and computational feasibility for all subsequent analyses.

Finally, to synthesize these findings, we calculated the intersection of programs identified by each setting with the 45 programs from our final model (Supp. Fig. S2). This analysis provides a global view of how each modeling choice affects the ability to discover biologically relevant signals. The results show that the single-cell approach is the most dissimilar, capturing the fewest of our model’s programs. Crucially, the setting lacking ligand competition also failed to identify a substantial fraction of programs, demonstrating that this component is vital for discovering more subtle and specific cellular interaction programs.

#### Settings compared

We generated sets of different numbers of K programs (from K=2 to K=100) using the 146 baseline MIBC samples as input. We compared 5 different settings:

1. The full model: containing the diffusion step and the competition step.
2. The full model with a modified input with random permutations across each slide. To do so, we permute for each sample the positions of the spots’ measured transcriptomics expression. Note that those permutations are independently done for each transcript. Then, we compute again for each randomized permutation the LR interaction and a new set of randomized programs.
3. An ablated version of the model without the diffusion step.
4. An ablated version of the model without the competition step.
5. The Tensor-cell2cell^38^ model using single-cell data as input. To do so, we used the cell2cell python package (version 0.7.4). We processed and used the matched single-cell data corresponding to the 146 baseline Visium samples we used in the other settings. We applied standard processing, using SCTransform^96^ and log-CPM normalization. We notably used the “expression_product” as the “communication_score” and the “non_negative_cp_hals” as the factorization algorithm.

#### Decomposition metrics

We define different metrics to compare decompositions obtained with different values of k and/or different settings.

##### Reconstruction error

The normalized reconstruction error is defined as the difference between the input matrix X and its approximation, normalized by the norm of the input matrix, such that:

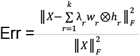

##### Cell type representation

Each obtained program can be characterized by the cell types involved, whether as senders or as receivers. We define two hierarchical levels of cell types. The first level corresponds to the triplet (Immune, Malignant, Stromal). The second level corresponds to the group (B cell, Mast, Granulocyte, MoMac, DC, T_NK, Epithelial, Endothelial, Fibroblast, Muscle, Malignant).

##### LR categories

The CellChatDB^91^ database provides annotations regarding the LR pairs mode of operation among: Cell-Cell Contact, Secreted Signaling, ECM Receptor and Non-protein Signaling. Considering a set of K programs, we thus characterize it by the number of unique LR pairs belonging to each of the four categories.

##### Subset of LRs characterizing a program

Due to the inherent sparsity of the output, we characterize a program based on subsets of LRs. Depending on the task:

- We preferably select the LRs with weights above a given threshold (0.1)
- If we need more accuracy while still reducing the vector’s dimension we select the LRs for which the sum of their squared weights is inferior or equal to a set threshold (0.99) when using it to compute a metric.

##### Number of LRs characterizing a program

Using the characteristic subset of LRs defined above, we define 4 possibilities to classify a program:

- “Single LR pair”: the subset contains only one LR pair.
- “Single Ligand across pairs”: the subset contains multiple LR pairs, but they all share the same ligand.
- “Single Receptor across pairs”: the subset contains multiple LR pairs, but they all share the same receptor.
- “Many LR pairs”: the subset contains multiple LR pairs.

##### Cosine similarity

Focusing on the vectors *h*_*r*_, representing the decomposition onto the LR pairs of a given program r, we can compare two programs from the same or from different decompositions through their cosine similarity, such that:

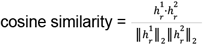

Note that we can compute this cosine similarity on the union of the two sets of characteristic LRs to reduce the vectors’ dimensions.

### Analysis tools

#### Sample-level descriptor

We define a sample-level descriptor for each program by using the sum of the program’s weights across the sample’s spots.

#### Association with molecular subtype

We assigned a molecular subtype to each sample using the consensusMIBC package^97^. The samples were classified into 6 molecular classes: Luminal Papillary (LumP), Luminal Non Specified (LumNS), Luminal Unstable (LumU), Stroma-rich, Basal/Squamous (Ba/Sq), Neuroendocrine-like (NE-like). No samples were classified as NE-like. We regrouped together the Luminal samples (LumP, LumNS and LumU). To statistically test the association of the program with the subtypes, we used pairwise Mann-Whitney U tests between the three classes using as input the sample-level descriptor of the program of interest.

#### Association with “T-cell aggregates” Status

We used a deep learning model trained to predict presence or absence of T-cell aggregates on matched H&E slides for every Visium sample^98^. For each slide and each spot (55um diameter), we find the H&E square tile of size 112 x 112 um. We run the model on that tile and obtain a score, which we binarize using a threshold of 0.5, leading to a classification of presence or absence of T-cell aggregates in each spot.

We then aggregate the program of interest’s spots’ weight for both categories and compare the distribution using a Mann-Withney U test at the cohort level.

#### Association with tumor regions

While noting that all Visium samples used were located in the Cancer Core, we define 3 regions within them: tumor islets, tumor islets edge and stroma. To do so, considering a Visium slide, we first define which spot contains tumor by using a threshold (0.2) on the normalized malignant cell type fraction. We then refine this tumor map to delineate cohesive tumor regions (the “islets”). The refinement process smoothed the tumor boundaries, connected nearby tumor spots, and filled any internal gaps. To ensure robustness, small, isolated tumor spots were filtered out, retaining only the most significant tumor areas. An “invasive margin” was then established by defining a boundary of a fixed width around the perimeter of these refined tumor islets. Based on this, every spot in the tissue was classified into one of three final regions: the Tumor Islet (the tumor’s core), the TI Edge (the invasive margin), or the Stroma (the surrounding non-tumoral tissue).

Similarly to the T-cell aggregates status, we computed the difference of the program of interest’s spots’ weights for both categories by using pairwise Mann-Whitney U tests.

#### Association with Transcription Factors activity

We first estimate the transcription factor (TF) activity within each spot of each Visium slide. To do so we use as reference the OMNIPATH^99^ dataset as prior knowledge and the decoupleR library. We use the Univariate Linear Model (ulm) method, which fits a linear model to predict the observed gene expression based on the TF’s TF-Gene interaction weight. The resulting score is the t-value of the slope obtained.

Once we have estimated the TF activity for a given sample, we evaluate its association with each of our programs using liana+’s bivariate local spatial metric method. It computes locally the association of a chosen metric (in our case the cosine similarity) between two variables, while weighting it with a radial decreasing function.

We aggregate this association across all samples and apply a multiple testing correction using the Benjamini-Hochberg procedure.

#### Association with gene signatures

Similarly to the TF activity inference, we are using Univariate Linear Model but using constructed networks corresponding to weighted genes in signatures of interest to quantify the activity of each signature across Visium spots

#### Association with Neo-Adjuvant Chemotherapy Response

A subset of the patients from the MIBC were treated with Neo-Adjuvant Chemotherapy. We have the treatment response information for 34 of those patients. We computed the association of the program of interest’s sample level descriptor with this information using a Mann-Whitney U test.

#### Tumor subpopulations analysis

One can estimate whether the programs are specific to a specific tumor subset.

First the snRNA-seq was used to identify tumor cells with different transcriptomics profiles.

Then these different groups of cells were mapped to the spatial transcriptomics data through deconvolution. In order to refine the arbitrarily stratified populations in snRNAseq (with a Louvain resolution of 0.4) we calculated the spatial correlation (Pearson, spotwise) per sample and grouped each pair of population with a person r > 0.5. Then, spatial association between these cancer cell populations and the different programs was performed with pearson and cosine liana local association.

#### P7 association with tumor fraction and CNV

To evaluate how Program 7 (P7), a tumor-associated communication program, relates to tumor fraction and chromosomal instability, we performed a mixed-effects regression analysis. First for the quantification of CNV_score, we used the inferCNVpy (python implementation of InferCNV^100^) to compute the CNV score, using the reference cell types from non-malignant populations (excluding “Malignant”, “Doublet”, “Other”, and “Unknown” cell types), and the following parameters: window size of 200 genes, step size of 10 genes, Leiden clustering resolution of 0.4, and using log1p CPM normalized counts. The analysis was performed on single-cell data from three cohorts with cells filtered to have at least 400 genes and fewer than 10,000 counts to ensure data quality. CNV_score was generated for each spot using the compute_cnv_score function.

All continuous variables (P7 activity, tumor fraction, and CNV score) were Z-score standardized to facilitate interpretation as standardized effect sizes. We fitted linear mixed-effects models with random intercepts and slopes grouped by patient (n=146 patients, 544,886 spatial spots), allowing both baseline P7 activity and the effects of tumor fraction and CNV score to vary across patients. This approach accounts for patient-level clustering and heterogeneity. Two models were compared: (1) a main effects model with tumor fraction and CNV score as predictors, and (2) an interaction model additionally including the Tumor_Fraction × CNV_score interaction term. Models were fitted using REML estimation with the Powell optimization algorithm. The random slopes model provided substantially better fit than random intercepts only (ΔLog-Likelihood = +111,811), revealing significant between-patient heterogeneity in CNV effects (variance=0.128) and tumor fraction effects (variance=0.072). The interaction model was selected as the final model. Although original variables showed moderate skewness, transformations degraded model fit and were not applied given the large sample size ensuring valid asymptotic inference via the Central Limit Theorem. Analyses were performed using Python 3.11 with statsmodels.

#### Mutation calling and identification of gene alterations associated with programs

Whole-exome sequencing (WES) data were generated as part of the MOSAIC Consortium project; details of the protocol and analysis are described in the associated publication^42^. Single-nucleotide variants (SNVs) and small insertions/deletions (indels) were annotated with the Ensembl Variant Effect Predictor (VEP) to identify affected genes and transcripts, as well as predicted variant consequences (e.g., nonsense, missense, frameshift).

Because MOSAIC WES data were derived from tumor samples only (without matched normal tissue), variants were classified as germline if present in gnomAD v4.0 with a population frequency above 0.0001 (based on the AF_grpmax metric, representing the maximum allele frequency across ancestry groups); otherwise, they were considered somatic.

Potentially oncogenic alterations were identified using the following criteria:

- Tumor suppressor genes (TSGs): High-impact variants (e.g., nonsense) in known TSGs from the IntOGen database. TSG classifications were pan-cancer, not cancer-type specific.
- Cancer hotspots: SNVs in protein-coding genes that produced previously reported amino acid substitutions at known cancer hotspots (from the CancerHotspots^101,102^ database and Hess et al. work^101,102^).

For copy-number variation (CNV) analysis, gene-level deletions and duplications were called. A CNV was considered to affect a gene if it overlapped at least one exon. Only high-quality CNVs (PASS in the VCF FILTER field) in protein-coding genes were retained. Genes impacted by CNVs were evaluated against the same TSG and oncogene sets used for mutation analysis.

Finally, we focused on genes annotated in the EpiFactors datababase^103^. This yielded a gene-by-sample matrix of alteration status. Based on this dataset and patient-level activity in P7, we calculated per-gene Mann-Whitney p-values, followed by multiple-testing correction using the Benjamini-Hochberg false discovery rate (FDR) method.

## Acknowledgments

We are grateful to patients, physicians, nurses and research assistants involved in the study. This study makes use of a NSCLC, BR, OV, MESO, DLBCL and GBM cohort of the MOSAIC consortium (Owkin; Charité –Universitätsmedizin Berlin (DE); Lausanne University Hospital - CHUV (CH); Universitätsklinikum Erlangen (DE); Institut Gustave Roussy (FR); University of Pittsburgh (USA)), a non-interventional clinical trial registered under NCT06625203.

