## Supplementary Figures for "Deciphering Cellular Ecosystems Driving Tumor Progression and Immune Escape from Spatial Transcriptomics and Single-Cell with COMPOTES"

#### Figures Index

| Chapter | Figure Nb | Title |
|---|---|---|
| An extension of existing methods to large scale spatial transcriptomics datasets | Supplementary Figure 1 | Ablation study - Metrics comparison for the compared settings across a fixed number of programs K ranging from 2 to 100. |
| An extension of existing methods to large scale spatial transcriptomics datasets | Supplementary Figure 2 | Ablation study - Comparison to the 45 programs. |
| Program 15 identifies a stalled or exhausted anti-tumor immunity | Supplementary Figure 3 | Description of the Program 15 |
| Program 15 identifies a stalled or exhausted anti-tumor immunity | Supplementary Figure 4 | Association with spot level features. |
| Deep dive into the program 7: A KMT2D-Linked Proliferative Signature Predicts Chemosensitivity in Early-Stage Bladder Cancer | Supplementary Figure 5 | Spot-level association of the 45 programs with different features, ranked by the average cosine similarity across all samples. |
| Deep dive into the program 7: A KMT2D-Linked Proliferative Signature Predicts Chemosensitivity in Early-Stage Bladder Cancer | Supplementary Figure 6 | Program 7 association with molecular features. |

### Deciphering Cellular Ecosystems Driving Tumor Progression and Immune Escape from Spatial Transcriptomics and Single-Cell with COMPOTES

#### Supplementary Figures: S1-6

### Figure S1. Ablation study - Metrics comparison for the compared settings across a fixed number of programs K ranging from 2 to 100.

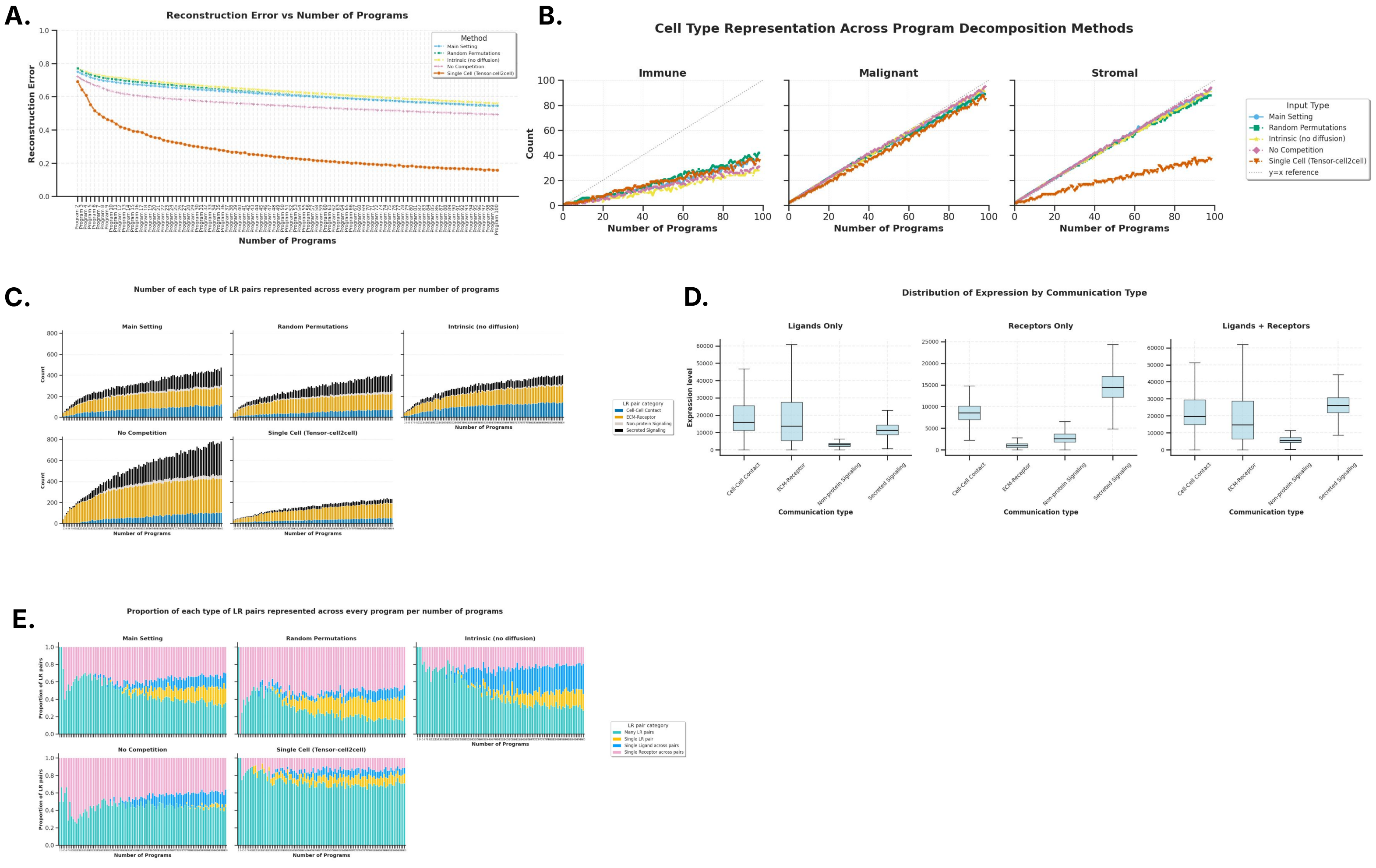


**Figure S1: Ablation study - Metrics comparison for the compared settings across a fixed number of programs K ranging from 2 to 100. A.** The evolution of the reconstruction error (see methods) **B.** The evolution of the number of programs involving cell types from one of the three categories: Immune, Malignant and Stromal cell types. **C.** The evolution of the number of LR pairs of each of the 4 categories describing the LR interaction mode, as described in the CellChat database: Cell-Cell contact, ECM Receptor, Non-protein Signaling and Secreted Signaling. We report the total number of the LR pairs from each of those categories across all programs for each number of programs K. **D.** Distribution of the expression values of the Ligands' and Receptors' genes in the L/R pairs input data, separated into the 4 categories described in the CellChat database. **E.**The evolution of the proportion of programs described by different categories of LR pairs. Either described by multiple LR pairs; by LR pairs sharing the same ligand; by LR pairs sharing the same receptors; or by a single LR pair.

### Figure S2. Ablation study - Comparison to the 45 programs.

A.

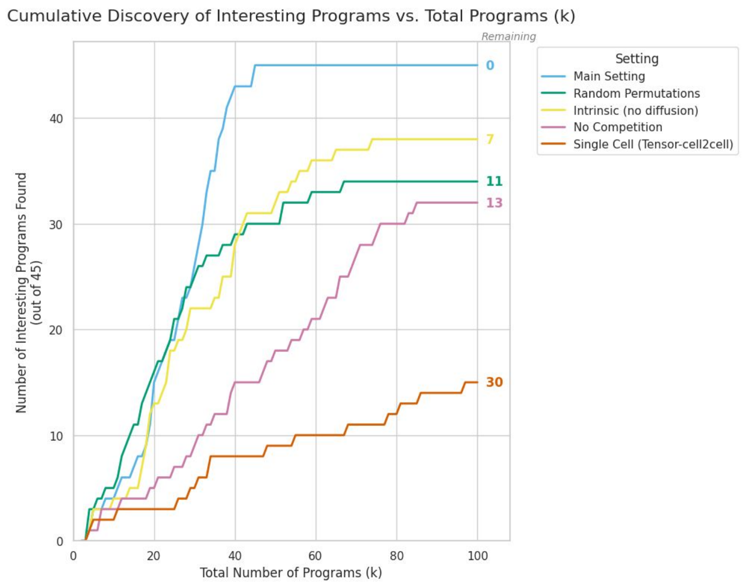


B.

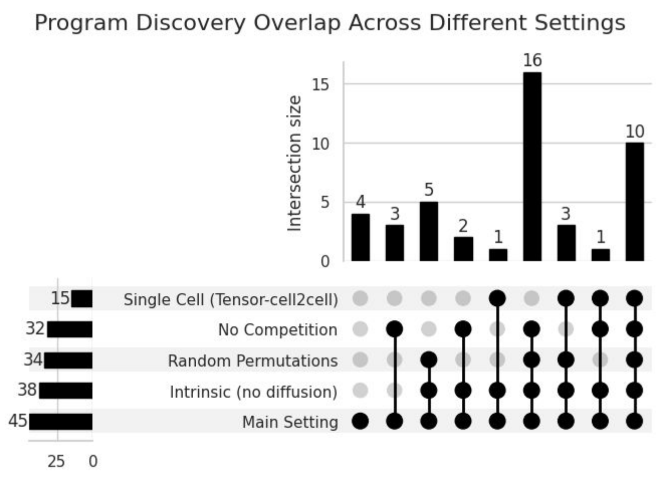


C.

| | N factors | N immune | Cell-Cell Contact | ECM Receptor | Secreted Signaling | Non-protein Signaling | Many LR | Single R | Single L | Single LR |
|---|---|---|---|---|---|---|---|---|---|---|
| Found in all | 10 | 3 | 6 | 3 | 1 | 0 | 3 | 5 | 1 | 1 |
| Found in all except single-cell | 16 | 11 | 9 | 9 | 6 | 2 | 8 | 14 | 2 | 2 |
| Found only in competition | 4 | 0 | 2 | 1 | 1 | 0 | 4 | 0 | 0 | 0 |
| Not found in intrinsic | 7 | 1 | 2 | 1 | 4 | 0 | 7 | 0 | 0 | 0 |
| Not found with random perm. | 11 | 3 | 4 | 1 | 6 | 0 | 11 | 0 | 0 | 0 |
| Not found without competition | 13 | 2 | 2 | 8 | 3 | 0 | 7 | 5 | 1 | 0 |
| Not found in single-cell | 30 | 10 | 7 | 11 | 10 | 2 | 16 | 11 | 2 | 1 |

**Figure S2: Ablation study - Comparison to the 45 programs**. For each setting, for a varying number of programs K ranging from 2 to 100, for each of the 45 programs of interest obtained with the main setting, we measured the first sighting of this program. **A.** The cumulative number of programs found at different numbers K for each setting. **B.** Analysis of which of the 45 programs were commonly found across which combinations of which settings. **C.** The analysis of the categories of programs that were found or not found in different settings or combinations of settings.

#### Figure S3. Description of the Program 15

*Figure S3 is on the next page*

**Figure S3: Description of the Program 15 A.** Top LR pairs of the program (weight ≥ 0.1). **B.** For each of the top LR pairs, mean expression and percentage of cell types expressing each ligand (top) and receptor (bottom) in single-cell data. **C.** Cellular network of Program 15 indicating the oriented weight of communication for the LR-pairs of this factor between each pair of cell types. The outer annotation color indicates the cell status (sender, receiver or both) while the inner color corresponds to each cell type to facilitate visualisation of links with regard to the sender cell type. **D.** Association of Program 15 with luminal and basal molecular subtypes at the patient level. **E.** Average activity of Program 15 expression across every sample's spots, stratified by the region where the spot is located (Tumor Islet, Tumor Islet's edge and Stroma). **F.** Distribution of Program 15 expression across every sample's spots, divided by whether a TLS was detected in that spot by the TLS Detect model. **G.** Example of one of the samples. The following are shown across spots: expression of Program 15; detection of TLS by the TLS Detect model; classification of the spots in the three regions defined; normalized deconvolution fractions of 6 cell types: malignant, fibroblast, natural killer t cells, monocytes/macrophages, dendritic cells and B cells.

### Supplementary Figure S3 (program 15 deep dive)

**A.**

| Ligand | Receptor | Weight |
|---|---|---|
| IL16 | CD4 | 0.2403 |
| C3 | C3AR1 | 0.2294 |
| CSF1 | CSF1R | 0.2174 |
| ANXA1 | FPR3 | 0.217 |
| C3 | ITGAX:ITGB2 | 0.2005 |
| PECAM1 | CD38 | 0.1664 |
| SEMA4D | PLXNB2 | 0.162 |
| LGALS9 | P4HB | 0.153 |
| ANXA1 | FPR1 | 0.1508 |
| GZMA | PARD3 | 0.1461 |
| NECTIN2 | TIGIT | 0.1389 |
| CD55 | ADGRE5 | 0.1383 |
| LGALS9 | HAVCR2 | 0.1383 |
| LGALS9 | CD44 | 0.1266 |
| APOE | TREM2:TYROBP | 0.1229 |
| CCL5 | CCR1 | 0.1202 |
| SELL | PODXL | 0.1157 |
| TNFSF14 | TNFRSF14 | 0.1145 |
| FASLG | FAS | 0.1139 |
| SELPLG | SELL | 0.1121 |
| THY1 | ADGRE5 | 0.11 |
| GAS6 | AXL | 0.1042 |
| CCL5 | CCR5 | 0.1038 |
| SEMA4D | CD72 | 0.1023 |

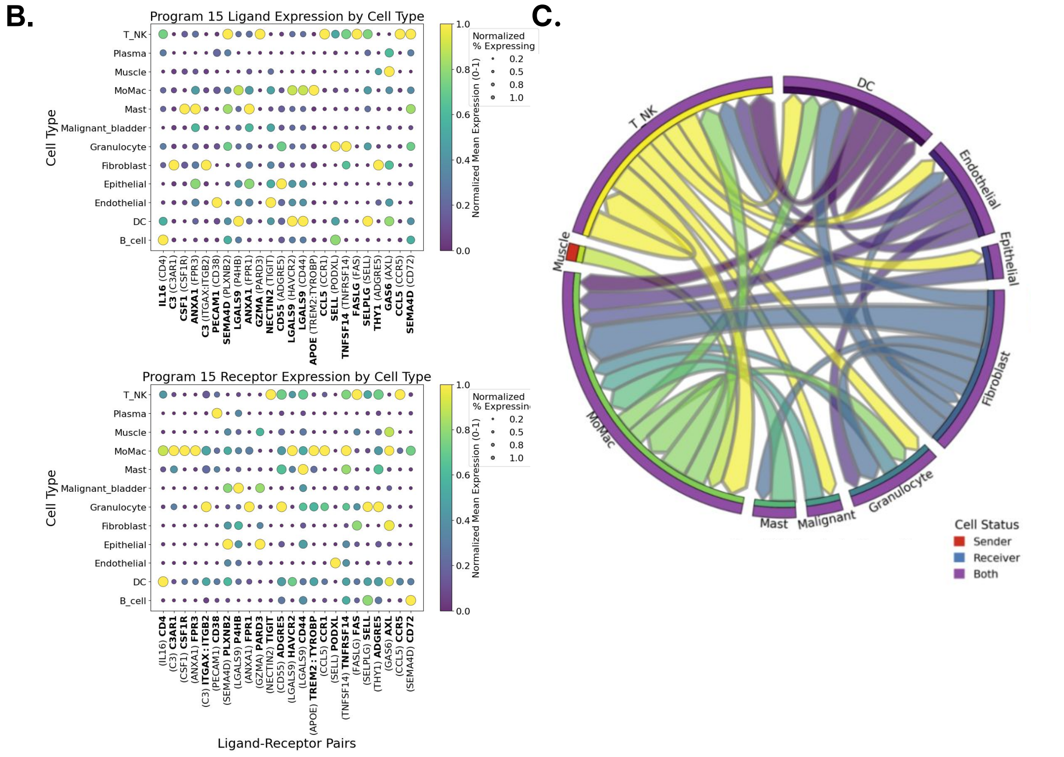


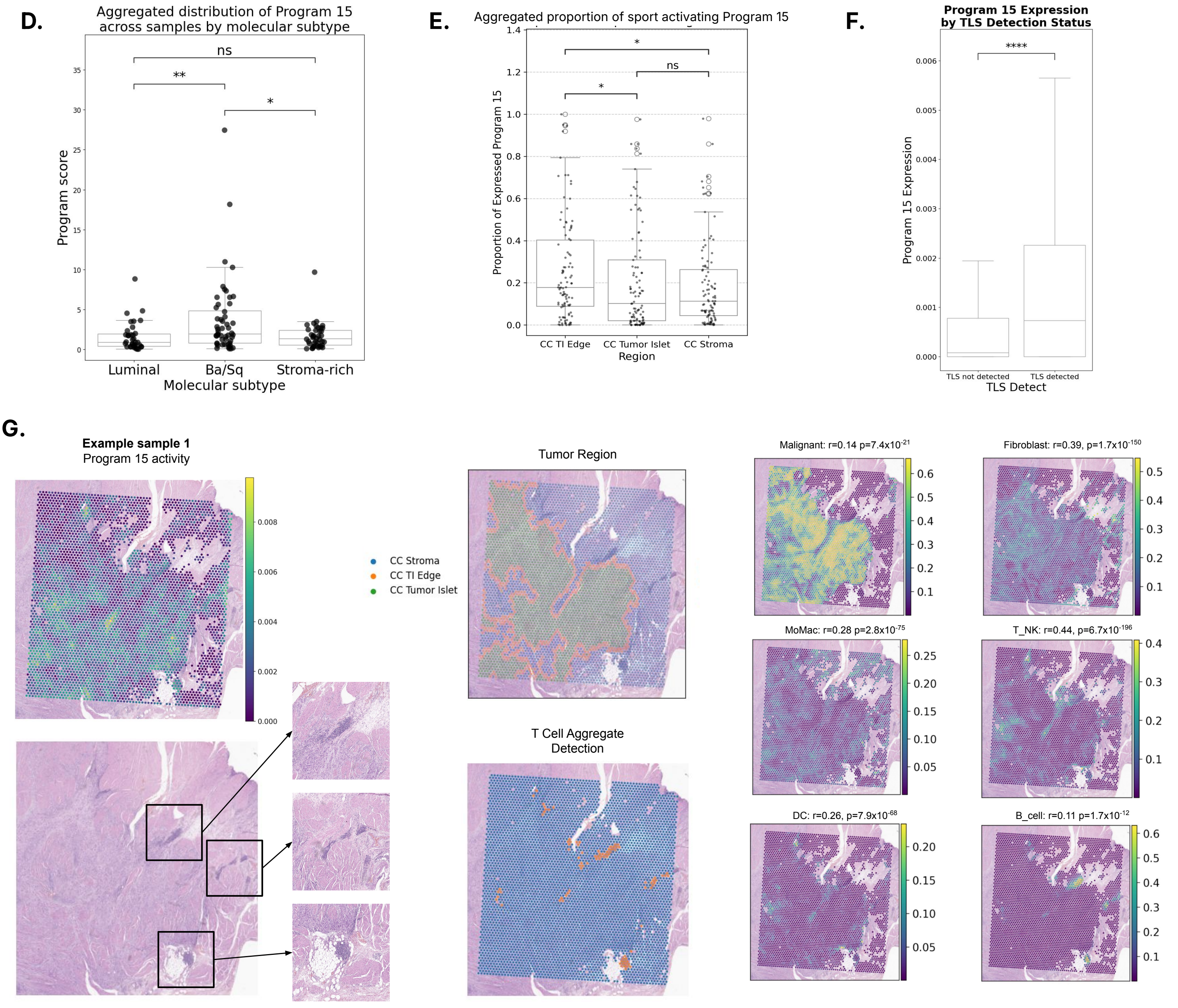

#### Figure S4. Association with spot level features.

*Figure S4 is on the next page*

**Figure S4: Association with spot level features.** Spot-level association of the 45 programs with different features, ranked by the average cosine similarity across all samples. **A.** Cell type deconvolution fractions at a first level of granularity. **B.** Cell type deconvolution fractions at a second level of granularity. **C.** Progeny pathways. **D.** Gene sets of interest from the literature.

Supplementary Figure S4 (program 15 associations)

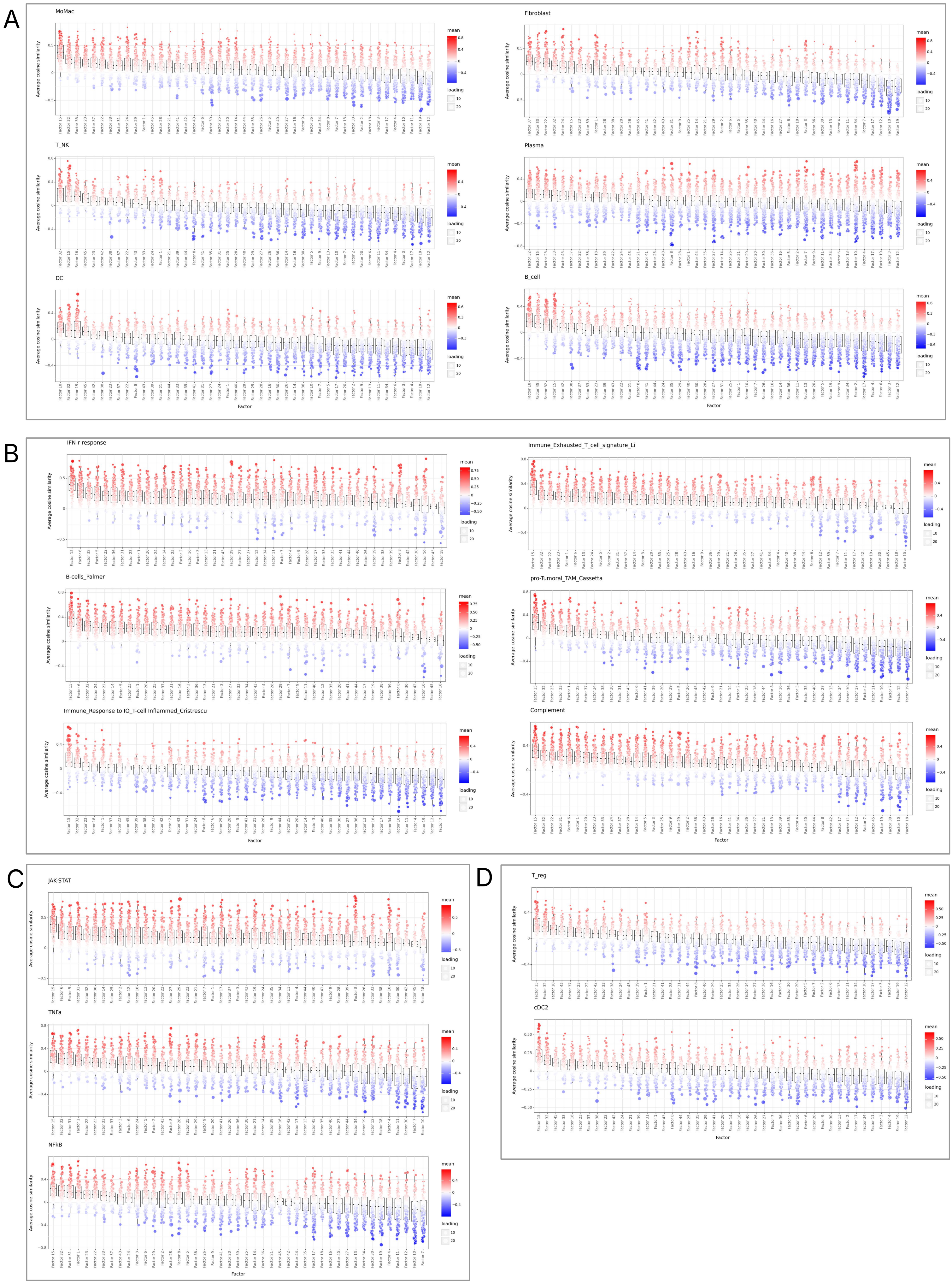

**Figure S5. Spot-level association of the 45 programs with different features, ranked by the average cosine similarity across all samples.**

*Figure S5 is on the next page*

**Figure S5: Spot-level association of the 45 programs with different features, ranked by the average cosine similarity across all samples. A.** Cell type deconvolution fractions at a first level of granularity. **B.** Progeny pathways. **C.** Gene sets of interest from the literature focusing on oncology pathways. **D.** Gene sets of interest from the literature focusing on immune pathways. **E.** Gene sets of interest from the literature focusing on response to treatment.

### Supplementary Figure S5 (program 7 associations)

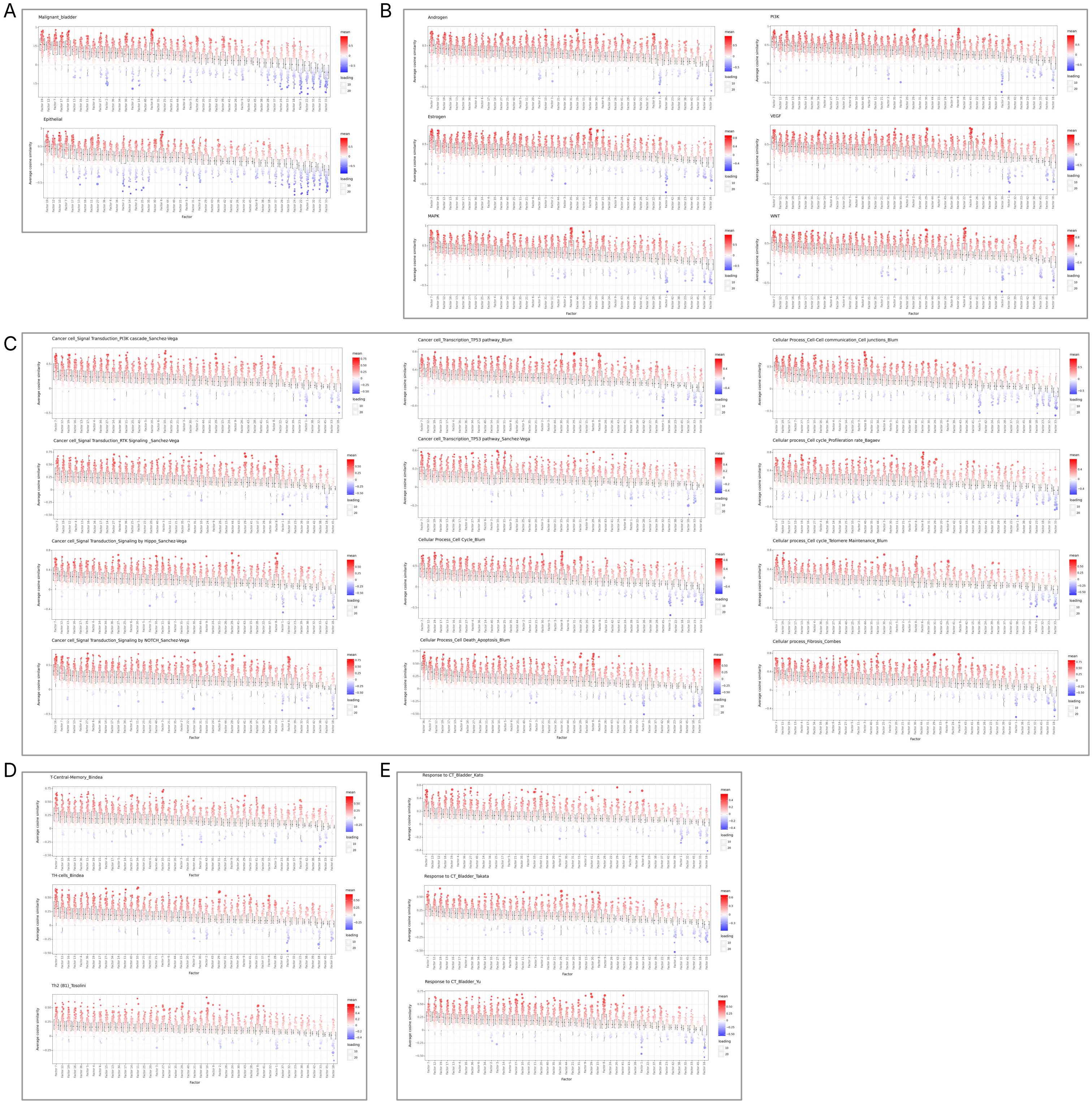

#### Figure S6. Program 7 association with molecular features.

*Figure S6 is on the next page*

**Figure S6: Program 7 association with molecular features. A.** Average activity of Program 15 expression across every sample's spots, stratified by the region where the spot is located (Tumor Islet, Tumor Islet's edge and Stroma). **C.** Oncoplot of TSG or Oncogenes with an alteration frequency above 20% in the cohort and belonging to the EpiFactors database. Color indicates the type of alteration in these genes for each patient. The number below the gene name corresponds to the corrected p-value (Mann-Whitney test with FDR-BH correction) testing if the activity of Program 7 is the same in the mutated and non-mutated groups. **C.** Bottom left heatmaps indicate an example of a sample with strong P7 activity in a region annotated as Epithelial. Histopathology zoom in the associated region (black square) indicate cells in a dysplastic state with pre-tumoral features. **D.** Results of a Mixed Effect Model with or without interaction between CNV_score and deconvolution malignant fraction to predict P7 activity across visium spots.

### Supplementary Figure S6 (cancer subpopulation association with P7)

**A.**

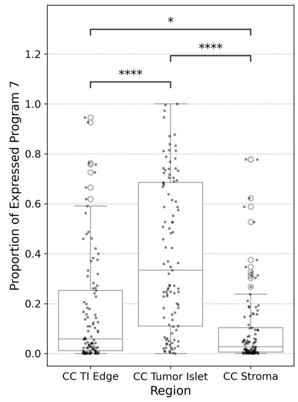


**B.**

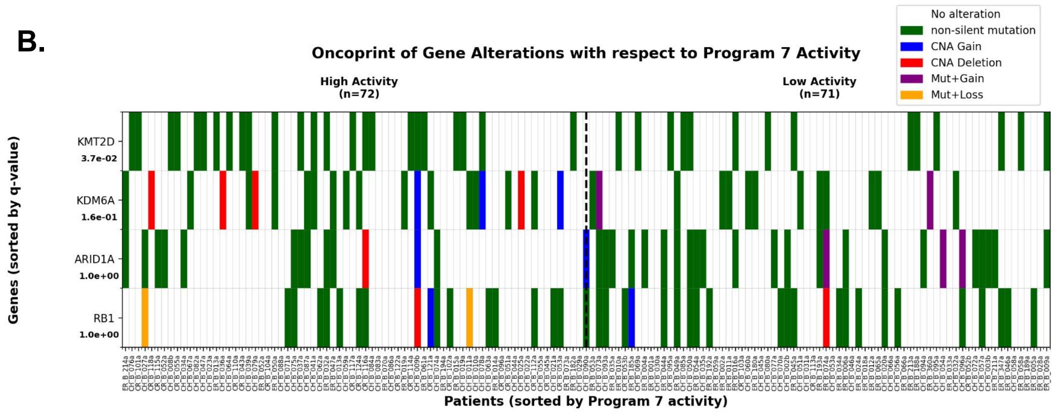


**C.**

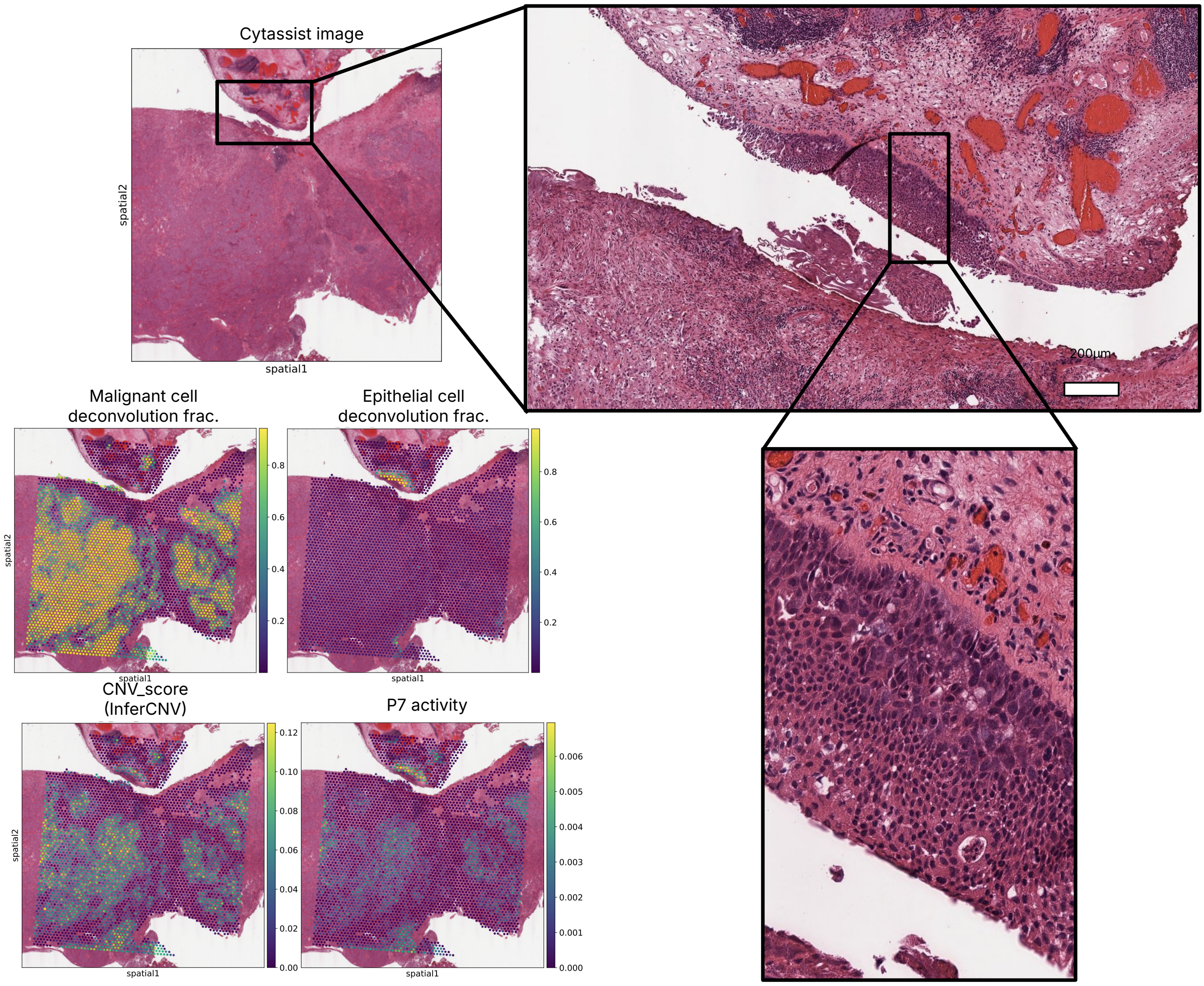


**D.** Estimates are fixed-effect coefficients from linear MXE.
Dependent variable: Program 7 activity; Random effect: Sample ID

| Model | Predictor | Estimate (β) | z | p-value |
|---|---|---|---|---|
| Without interaction | Tumor fraction | 0.060 | 2.71 | 0.007 |
| | CNV score | 0.430 | 14.50 | $<1x10^{-47}$ |
| With interaction | Tumor fraction | 0.066 | 2.97 | 0.003 |
| | CNV score | 0.420 | 14.27 | $<1x10^{-45}$ |
| | Tumor fraction × CNV score | 0.015 | 11.90 | $<1x10^{-32}$ |
